# Chiropterans are a hotspot for horizontal transfer of DNA transposons in Mammalia

**DOI:** 10.1101/2023.03.23.533946

**Authors:** Nicole S Paulat, Jessica M Storer, Diana D Moreno-Santillán, Austin B Osmanski, Kevin AM Sullivan, Jenna R Grimshaw, Jennifer Korstian, Michaela Halsey, Carlos J Garcia, Claudia Crookshanks, Jaquelyn Roberts, Arian FA Smit, Robert Hubley, Jeb Rosen, Emma C Teeling, Sonja C Vernes, Eugene Myers, Martin Pippel, Thomas Brown, Michael Hiller, Zoonomia Consortium, Danny Rojas, Liliana M Dávalos, Kerstin Lindblad-Toh, Elinor K Karlsson, David A Ray

**Affiliations:** Department of Biological Sciences, Texas Tech University; Lubbock, TX 79409, USA; Institute for Systems Biology; Seattle, WA 98109, USA; School of Biology and Environmental Science, University College Dublin; Belfield, Dublin 4, Ireland; Neurogenetics of Vocal Communication Group, Max Planck Institute for Psycholinguistics; 6525 XD, Nijmegen, The Netherlands; Donders Institute for Brain, Cognition and Behaviour; 6525 AJ, Nijmegen, The Netherlands; School of Biology, The University of St Andrews; Fife KY16 9ST, UK; Max Planck Institute of Molecular Cell Biology and Genetics; 01307, Dresden, Germany; LOEWE Centre for Translational Biodiversity Genomics; 60325, Frankfurt, Germany; Department of Natural Sciences and Mathematics, Pontificia Universidad Javeriana Cali, Valle del Cauca, Colombia; Department of Ecology and Evolution, Stony Brook University; Stony Brook, NY 11790, USA; Consortium for Inter-Disciplinary Environmental Research, Stony Brook University; Stony Brook, NY 11790, USA; Science for Life Laboratory, Department of Medical Biochemistry and Microbiology, Uppsala University; Uppsala, 751 32, Sweden; Broad Institute of MIT and Harvard; Cambridge, MA 02139, USA; Program in Bioinformatics and Integrative Biology, UMass Chan Medical School; Worcester, MA 01605, USA; Program in Molecular Medicine, UMass Chan Medical School; Worcester, MA 01605, USA

## Abstract

Horizontal transfer of transposable elements is an important mechanism contributing to genetic diversity and innovation. Bats (order Chiroptera) have repeatedly been shown to experience horizontal transfer of transposable elements at what appears to be a high rate compared to other mammals. We investigated the occurrence of horizontally transferred DNA transposons involving bats. We found over 200 putative horizontally transferred elements within bats; sixteen transposons were shared across distantly related mammalian clades and two other elements were shared with a fish and two lizard species. Our results indicate that bats are a hotspot for horizontal transfer of DNA transposons. These events broadly coincide with the diversification of several bat clades, supporting the hypothesis that DNA transposon invasions have contributed to genetic diversification of bats.

## Introduction

Transposable elements (TEs), DNA fragments that can mobilize within and across genomes, comprise most horizontally transferred (HT) genetic material in eukaryotes (Wallau, et al. 2012; El Baidouri, et al. 2014). Although viruses are prime candidates as TE vectors (Gilbert, et al. 2010; Thomas, et al. 2010; Gilbert, et al. 2014; Gilbert, et al. 2016), the exact mechanisms of how TEs are transferred and invade the germline of eukaryotes are unclear. Nevertheless, horizontal transfer of transposable elements (HTT) into naïve genomes can allow TEs to successfully invade and propagate before the host can effectively silence the invaders with anti-TE defenses (Schaack, et al. 2010; Kofler, et al. 2018). Class II elements (DNA transposons and rolling-circle (RC) elements), particularly Tc-Mariner transposons, are overrepresented in eukaryote HT events compared to Class I elements (retrotransposons) (Peccoud, et al. 2017; Zhang, et al. 2020), likely due to differences in mobilization mechanisms allowing easier transmission (Lampe, et al. 1996; Silva, et al. 2004; Gilbert, et al. 2016; Gilbert and Feschotte 2018; Palazzo, et al. 2019).

The activity and repetitive nature of TEs have shaped genome structure and phenotypes in diverse lineages, by increasing TE copy number, introducing genetic diversity, altering regulatory networks, and promoting shuffling of exons and by introducing TE domains that can be coopted by the host genome (Feschotte and Pritham 2007; Feschotte 2008; Cordaux and Batzer 2009; Schaack, et al. 2010; Casacuberta and González 2013; Thomas, et al. 2014; Grabundzija, et al. 2016; Zhang, et al. 2019; Cosby, et al. 2021). Yet the magnitude of influence on genome evolution in mammals is unclear, as previous studies were limited by relatively few mammal genome assemblies and TE datasets. High sequence similarity among observed DNA transposons and relatively recent divergence of many mammal lineages make it difficult to parse HTT versus vertical inheritance (Gilbert, et al. 2010; Novick, et al. 2010; Zhang, et al. 2020). Recent publication of many genome assemblies from diverse species has resolved at least one of these problems (Genereux, et al. 2020; Jebb, et al. 2020; Rhie, et al. 2021; Threlfall and Blaxter 2021), creating an opportunity to determine the extent of HTT.

Mammalian genomes are of considerable interest due to their propensity for relatively low TE diversity compared to most other vertebrates (Furano, et al. 2004; Chalopin, et al. 2015; Sotero-Caio, et al. 2017), making HTT events more easily identifiable. While typically 20-50% of mammalian genomes are TE-derived, much of this is from retrotransposons (Chalopin, et al. 2015; Sotero-Caio, et al. 2017); most mammals have experienced little to no DNA transposon accumulation in the last 40 My (Pace and Feschotte 2007; Sotero-Caio, et al. 2017). A major exception to this observation is the order Chiroptera, especially members of the family Vespertilionidae, which are well-known for having unusually diverse TE repertoires, and experiencing several recent, independent DNA transposon invasions (Pritham and Feschotte 2007; Ray, et al. 2007; Ray, et al. 2008; Thomas, et al. 2011; Pagán, et al. 2012; Mitra, et al. 2013; Ray, et al. 2015; Platt, et al. 2016). While the impacts of these DNA transposon invasions are not fully understood, they offer a large pool of genetic variation that may contribute to rapid genome evolution in bats. Several studies have shown TE-driven exon shuffling and transposase cooption have impacted bat evolution (Pritham and Feschotte 2007; Thomas, et al. 2014; Grabundzija, et al. 2016; Cosby, et al. 2021). Indeed, a fair number of DNA-transposon derived genes are found in mammal and vertebrate lineages with a variety of functions including, but not limited to transcription, chromosome structure, and immunity (reviewed in Feschotte and Pritham 2007).

Bats are the second largest order of mammals (n∼1426), exhibiting some of the most unique mammalian phenotypes (e.g., flight, laryngeal echolocation, extended longevity, tolerant immunity) and inhabiting multiple ecological niches (*24*). This phenotypic diversity along with their unusual diversity of younger TEs led us to investigate HT of DNA transposons involving bats. In addition to the broad array of mammalian genomes from the Zoonomia Project (Genereux, et al. 2020), several bat genome assemblies have been produced by the Bat1K Project (Teeling, et al. 2018; Jebb, et al. 2020). Combined, this genomic data includes thirty-seven bat species from 11 families and 28 genera spanning the two major chiropteran clades, Yinpterochiroptera and Yangochiroptera (Teeling, et al. 2005; Amador, et al. 2018). We analyzed TE accumulation patterns across Chiroptera and leveraged TE curation data from 251 mammal assemblies to perform a large-scale analysis of recent HT of DNA transposons involving bats. Our findings highlight TE-based diversity within bats and suggest that, in a radical departure from other eutherian mammals, Chiroptera is a hotspot for HTT.

## Results

### More Recent, Substantial DNA Transposon Accumulation in Bats

We used a curated *de novo* TE library to annotate TE insertions in 250 eutherian mammalian species, including 37 bat species (Table S1) (Osmanski, et al. 2022; Christmas, et al. forthcoming). A general comparison of TE content among mammal assemblies is available elsewhere (Osmanski, et al. 2022). Rather than recapitulate that work in illustrating general distinctions between bats and non-bats, we chose eight representative eutherians as our outgroup taxa. Of the eight, four species were selected due to having the greatest accumulation of young (≤50 My) DNA transposons outside of bats: two tenrecs (*Echinops telfairi* and *Microgale talazaci*, Afrosoricida), and the Eastern mole and the Indochinese shrew (*Scalopus aquaticus* and *Crocidura indochinensis*, Eulipotyphla). The other four species along with the eulipotyphlans represent one of the five mammalian orders closely related to Chiroptera within Laurasiatheria (Foley, et al. 2022): horse (*Equus caballus*, Perissodactyla), cow (*Bos taurus*, Artiodactyla), pangolin (*Manis javanica*, Philodota), and domestic cat (*Felis catus*, Carnivora).

With regard to total TE content, bats generally resemble other mammals, with TEs composing 30-60% of the genome, with 15-30% from LINE elements, and the rest split among SINE, LTR, and DNA elements (Fig. 1a). The eight outgroup mammals are similar in proportions of different types of TEs, though the eulipotyphlans have slightly lower TE content overall, and *Bos taurus* harbors a relatively high proportion of LINEs (Osmanski, et al. 2022). The latter has been discussed previously and is due to an independent HT of RTE-like retrotransposons, Bov-B LINEs (Kordis and Gubensek 1998). Such variation in retrotransposon content is not unexpected among mammals (Sotero-Caio, et al. 2017; Platt, et al. 2018).

**Fig. 1:**
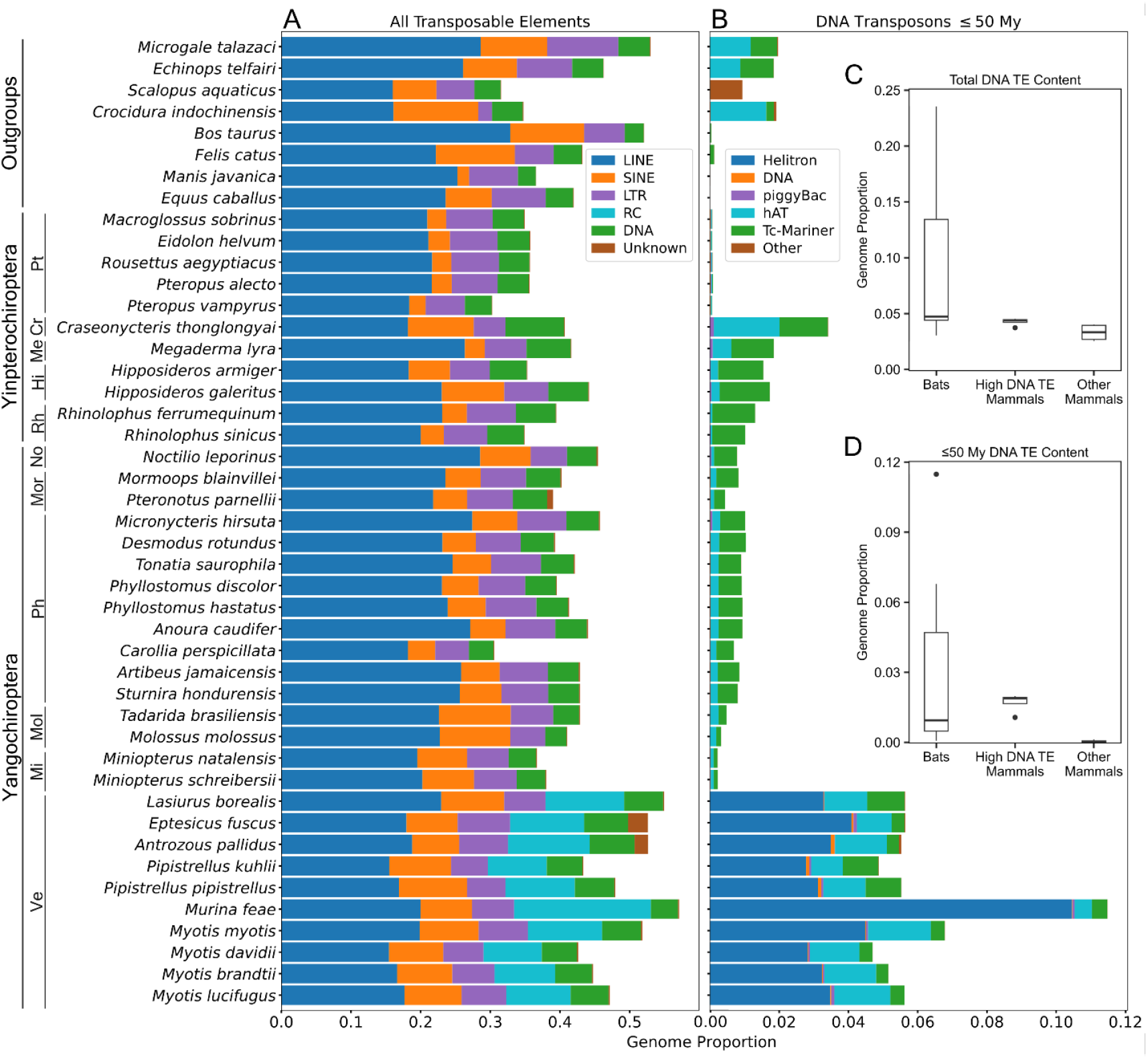
(**A**) Total transposable element accumulation, (**B**) DNA transposon accumulation within the last 50 My, and (**C and D**) box plots depicting ranges of total DNA transposon genome content in 37 chiropterans and 8 outgroup mammalians. High DNA TE Mammals are defined as described in the main text as *Echinops telfairi, Microgale talazaci, Scalopus aquaticus* and *Crocidura indochinensis*. Bat families are indicated by abbreviations left of species names and are as follows: Pt = Pteropodidae, Me = Megadermatidae, Cr = Craseonycteridae, Rh = Rhinolophidae, Hi = Hipposideridae, Ve = Vespertilionidae, Mi = Miniopteridae, Mol = Molossidae, No = Noctilionidae, Mor = Mormoopidae, Ph = Phyllostomidae.

However, there are several major differences between bats and non-bats. Most notable is the presence of generally higher total and more recent DNA transposon accumulation (Fig. 1b-d), mostly hAT and Tc-Mariner transposons, in many of the bat subclades and the obvious presence of substantial accumulation of RC elements in vespertilionid bats in the last 50 My (Fig. 1b). Substantial RC accumulation is not observed in yinpterochiropteran bats or outgroup species. Within the DNA transposon categories, vespertilionid bats also have higher hAT element accumulation than yinpterochiropteran lineages, except for the bumblebee bat (*Craseonycteris thonglongyai*) and the lesser false vampire bat (*Megaderma lyra*) (Fig. 1b). In comparison to non-bats, vespertilionid bats and *Craseonyteris thonglongyai* have higher young DNA transposon accumulation than all outgroup mammals, but the four high DNA TE mammals have greater amounts of young DNA transposons than most if not all other bats. However, the other four mammals, have less young DNA transposon accumulation than all bats expect pteropodids, and this low recent accumulation is more representative of eutherian mammals in general (Table S2) (Osmanski, et al. 2022).

### Temporal Class II Transposon Accumulation in Bats

To examine the temporal context of TE accumulation, we calculated each TE copy’s divergence from the TE consensus sequence and applied species-specific neutral mutation rates (Table S3) to assign insertion times to each insertion. To explore temporal variation in Class II accumulation among lineages, we visualized DNA/RC accumulation within the past ∼50 My in Fig. 2. This figure illustrates broad patterns of DNA transposon superfamily accumulation as it varies by bat family and patterns that are clearly lineage specific. Each superfamily comprises multiple, potentially lineage-specific subfamilies.

**Fig. 2.**
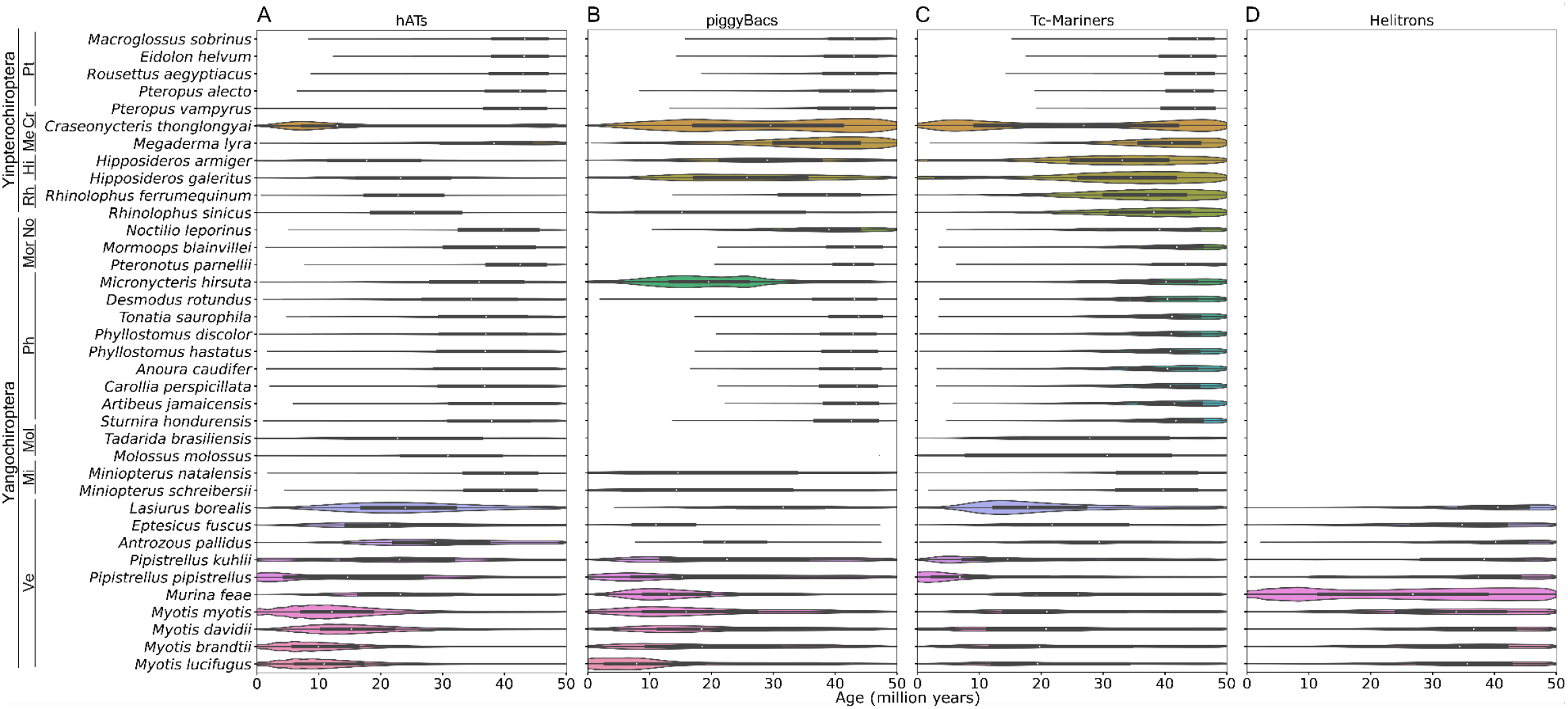
Violin plots of DNA transposon distributions by family in bats. Distributions of (**A**) hAT, (**B**) piggyBac, (**C**) Tc-Mariner, and (**D**) Helitron elements within the last 50 million years in 37 bat species. Species are arranged phylogenetically; bat families are indicated by abbreviations left of species names and are as follows: Pt = Pteropodidae, Me = Megadermatidae, Cr = Craseonycteridae, Rh = Rhinolophidae, Hi = Hipposideridae, Ve = Vespertilionidae, Mi = Miniopteridae, Mol = Molossidae, No = Noctilionidae, Mor = Mormoopidae, Ph = Phyllostomidae.

For example, vespertilionid bats (Yangochiroptera) show substantial hAT accumulation within the last 40 My, with *Myotis* species showing the highest hAT accumulation between 10-20 Mya, coinciding with species diverging between 10.9-18.2 Mya (Kumar, et al. 2022), while *Lasiurus borealis* appears to have experienced a slightly older peak of accumulation 20-35 Mya (Fig. 2a). The two available *Pipistrellus* species have experienced increased hAT accumulation within the last 5 My, well after the divergence of the two species ∼9.6-17.6 Mya (Kumar, et al. 2022). All vespertilionid bats show Helitron accumulation across the last 50 My, including ancestral accumulation, but *Murina feae* displays a surprisingly large amount, with accumulation peaks ∼10 and 40 Mya (Fig. 2d). Across other yangochiropterans, *Micronycteris hirsuta* stands out as experiencing a burst of piggyBac accumulation not apparent in other phyllostomids (Fig. 2b), otherwise phyllostomids show consistent patterns of ancestral Tc-Mariner accumulation 40-50 Mya and little else (Fig. 2). *Noctilio leporinus* shows high Tc-Mariner accumulation over the span of 25-50 Mya, with little accumulation more recently (Fig. 2c).

Yinpterochiropterans display similarly variable Class II accumulation (Fig. 2). Pteropodid bats display a uniform lack of substantial DNA transposon accumulation within the last 50 My, with little to no accumulation within the last 10 My (Fig. 2). This is consistent with previous observations of no substantial retrotransposon accumulation over approximately the same period (Cantrell, et al. 2008; Nikaido, et al. 2020). Other yinpterochiropterans show peaks of Tc-Mariner accumulation 35-40 Mya, and low-level accumulation of other DNA transposons. *Craseonycteris thonglongyai* and its closest relative in this study, *Megaderma lyra*, both have considerably higher piggyBac accumulation, and to a much lesser extent hAT accumulation than other yinpterochiropterans. However, *C. thonglongyai* also exhibits a striking increase of species-specific DNA transposon accumulation in the last 5-6 My, with a second peak of hAT, piggyBac, and Tc-Mariner accumulation (Fig. 2a-c).

### Many More HT Events in Bats Compared to Other Mammals

Lineage-specific TE subfamilies constitute much of the DNA and RC accumulation across bat lineages in the last 50 My, an observation consistent with previous studies (Pritham and Feschotte 2007; Ray, et al. 2007; Ray, et al. 2008; Thomas, et al. 2011; Pagán, et al. 2012; Mitra, et al. 2013; Zhuo, et al. 2013; Platt, et al. 2016). Unlike LINE retrotransposons, which tend to accumulate over long periods and exist as multiple lineages in genomes, diversifying into sometimes numerous subfamilies (Konkel, et al. 2010; Boissinot and Sookdeo 2016), DNA transposons are prone to inactivating internal deletions and tend to have shorter lifespans (Lohe, et al. 1995; Smit 1996; Feschotte and Pritham 2007; Muñoz-López and García-Pérez 2010; Gilbert and Feschotte 2018). As a result, recent accumulation of a wide variety of DNA transposons is intriguing and suggests possible external origins.

Historically, the criterion used to identify a potential HTT is the presence of a unique TE in a given genome and the corresponding absence from close relatives. While not always possible, confirming the presence of a highly similar element in the genome of a distant relative serves as strong confirmation of the HTT. An example is the presence of a piggyBac transposon, *piggyBac2_ML*, in the *Myotis lucifugus* genome, and a highly similar element, *piggyBac2_Mm*, in the genome of *Microcebus murinus*, a lemur (Pagan, et al. 2010). The concurrent absence of any similar elements in the genomes of other mammals strongly suggests horizontal movement from one lineage to the other via some, usually unknown, vector, such as a virus (Gilbert, et al. 2010; Thomas, et al. 2010; Gilbert, et al. 2014; Gilbert, et al. 2016; Gilbert and Feschotte 2018).

We investigated possible HT of bat DNA transposons across mammals and other eukaryotes using a broad-scale approach (Materials and Methods). We identified 221 putative HT DNA/RC transposons representing 229 HT events involving bats (Table S4, S5, S6). Tc-Mariner elements are well-known as frequent participants in HT (Peccoud, et al. 2017; Reiss, et al. 2019; Zhang, et al. 2020), and as expected, comprise over a third of putative HT events (n = 84, 36.7%). Elements from the hAT, piggyBac, and Helitron families make up the remaining 145 HT events (n = 64, 29, 52, respectively). BLAST searches indicated no copies of these putative HTTs in any available eukaryote assembly (other than the chiropteran assemblies from which it was originally detected) in all but 19 cases (see below). Previous studies (Wallau, et al. 2012; Melo and Wallau 2020) have also used searches of orthologous insertion sites in addition to BLAST to confirm patchy TE distributions of putative HTTs. However, the large number of mammal assemblies and putative HTTs precluded such a large number of additional searches. We therefore queried two outgroup species with high quality genome assemblies, *Bos taurus* and *Equus caballus*, in detail for orthologous TE copies of the 221 putative HTTs. These searches yielded zero full-length or partial matches. These results along with the lack of BLAST hits are consistent with horizontal transfer rather than prolonged vertical transmission.

Of the nineteen HTTs where a non-chiropteran match was identified by BLAST, sixteen elements involved other eutherian clades including Lemuriformes (12 TEs), Afrosoricida (6 TEs), Scandentia (1 TE), and Eulipotyphla (1 TE) (Table 1, Table S5, S7). These HTTs included ten hAT elements, five Tc-Mariner elements and two piggyBac elements. Two HTTs, *Mariner_Tbel* and *npiggy1_Mm*, were previously identified as horizontal transfers involving mammals. *Mariner_Tbel* was previously found in the tree shrew *Tupaia belangeri* (Oliveira, et al. 2012), consistent with our findings (Table S5, S7), as well as the European hedgehog *Erinaceus europaeus. npiggy1_Mm*, a non-autonomous piggyBac element previously identified as part of an HT event with its autonomous partner *piggyBac1_Mm* in the lemur *Microcebus murinus* (Pagan, et al. 2010). Zero orthologous HTT insertions were found between these mammals and bats indicating independent insertion events consistent with HT. A single autonomous hAT element, *OposCharlie2*, was found in a marsupial, *Monodelphis domestica*, consistent with previous HT studies (Gilbert, et al. 2010; Novick, et al. 2010). Only two elements were detected in non-mammals. An autonomous Tc-Mariner, *Mariner2_pKuh*, was found in an African reedfish, *Erpetoichthys calabaricus*, and the bat *Pipistrellus kuhlii* (52 and 327 copies, respectively (Table S7)), but not in the closely related *Pipistrellus pipistrellus*. This is consistent with the estimated age of the element, ∼2.2 My, which is younger than the divergence of the two pipistrelle species, ∼10 to 18 Mya (Kumar, et al. 2022). The element has high sequence conservation as well, with 99.74% identity between the two species’ consensus sequences (Fig. S1). The second element, a non-autonomous Tc-Mariner, *nMariner1_Lbo*, was identified in two lizard species, *Zootoca vivipara* and *Lacerta agilis*, as well as three vespertilionid bats (Table S7), with sequence conservation of >83% among all species, >90% excluding the single insertion in *Antrozous pallidus* (Fig. S2). Only five of the nineteen putative HTTs are autonomous. Our methods assumed that many possible autonomous HTTs have <90 annotated copies in bat genomes, possibly due to loss or degradation, but that the corresponding HT events are represented by these non-autonomous counterparts.

**Table 1.** Summary of putative horizontally transferred DNA transposons present in multiple eukaryote clades.

| TE Subfamily | TE Family | Consensus<br>Length (bp) | Number of Species Involved |  |  |  |  |  |  |  |
| --- | --- | --- | --- | --- | --- | --- | --- | --- | --- | --- |
|  |  |  | Polypteriformes | Squamata | Didelphimorphia | Afrosoricida | Scandentia | Lemuriformes | Eulipotyphla | Chiroptera |
| *CraTho-1.191* | hAT | 191 | 0 | 0 | 0 | 0 | 0 | 4 | 0 | 5 |
| *CraTho-2.327* | hAT | 213 | 0 | 0 | 0 | 1 | 0 | 0 | 0 | 3 |
| *EchTel-1.100* | hAT | 334 | 0 | 0 | 0 | 2 | 0 | 0 | 0 | 5 |
| *hAT-2N1_MM* | hAT | 198 | 0 | 0 | 0 | 1 | 0 | 4 | 1 | 7 |
| *MirCoq-4.2925* | hAT | 223 | 0 | 0 | 0 | 0 | 0 | 3 | 0 | 2 |
| *MurFea-1.231* | hAT | 230 | 0 | 0 | 0 | 1 | 0 | 3 | 0 | 2 |
| *MyoBra-4.2938* | hAT | 229 | 0 | 0 | 0 | 0 | 0 | 2 | 0 | 4 |
| *MyoBra-5.81* | hAT | 192 | 0 | 0 | 0 | 0 | 0 | 4 | 0 | 5 |
| *OposCharlie2* | hAT | 2996 | 0 | 0 | 1 | 0 | 0 | 0 | 0 | 2 |
| *SPIN_NA_9_Ml* | hAT | 311 | 0 | 0 | 0 | 2 | 0 | 0 | 0 | 5 |
| *SPIN_Og* | hAT | 2836 | 0 | 0 | 0 | 2 | 0 | 2 | 0 | 13 |
| *npiggy1_Mm* | piggyBac | 240 | 0 | 0 | 0 | 0 | 0 | 2 | 0 | 1 |
| *npiggyBac-2_EF* | piggyBac | 172 | 0 | 0 | 0 | 0 | 0 | 2 | 0 | 1 |
| *DNA2_pKuh* | Tc-Mariner | 152 | 0 | 0 | 0 | 0 | 0 | 1 | 0 | 4 |
| *Mariner2_pKuh* | Tc-Mariner | 2292 | 1 | 0 | 0 | 0 | 0 | 0 | 0 | 1 |
| *Mariner3_pKuh* | Tc-Mariner | 2282 | 0 | 0 | 0 | 0 | 0 | 1 | 0 | 2 |
| *Mariner_Tbel* | Tc-Mariner | 1283 | 0 | 0 | 0 | 0 | 2 | 0 | 0 | 3 |
| *nMar1_Rf* | Tc-Mariner | 236 | 0 | 0 | 0 | 0 | 0 | 2 | 0 | 6 |
| *nMariner1_Lbo* | Tc-Mariner | 184 | 0 | 2 | 0 | 0 | 0 | 0 | 0 | 3 |
| Total # Unique Species |  |  | 1 | 2 | 1 | 2 | 2 | 5 | 1 | 18 |

In contrast to the 229 HT events in bats, few possible HT events were identified in other mammals (detailed above and in Christmas et al. (forthcoming)). Of the six other orders with HT events, only Primates and Afrosoricida had more than five events (15 and 6, respectively). To compare HT events between the 37 bats and 213 other eutherian mammal species, we modeled the number of events by mammalian order (Table S8) using a negative binomial distribution and estimated HT means for both bats and non-bats. Although bats represent only one mammalian order, this point observation can be compared to the posterior distribution of the mean of HT events across eighteen other orders (equivalent to a one sample t-test for normal data). As there is only a single order to estimate the mean for bats, posterior distribution of these estimates overlap (Fig. 3). However, considering there is only a single point estimate of HT for bats, it does not overlap with the posterior mean of HT for all other mammalian orders. This demonstrates that there were many more HT events in bats than in other mammalian orders.

**Fig. 3.**
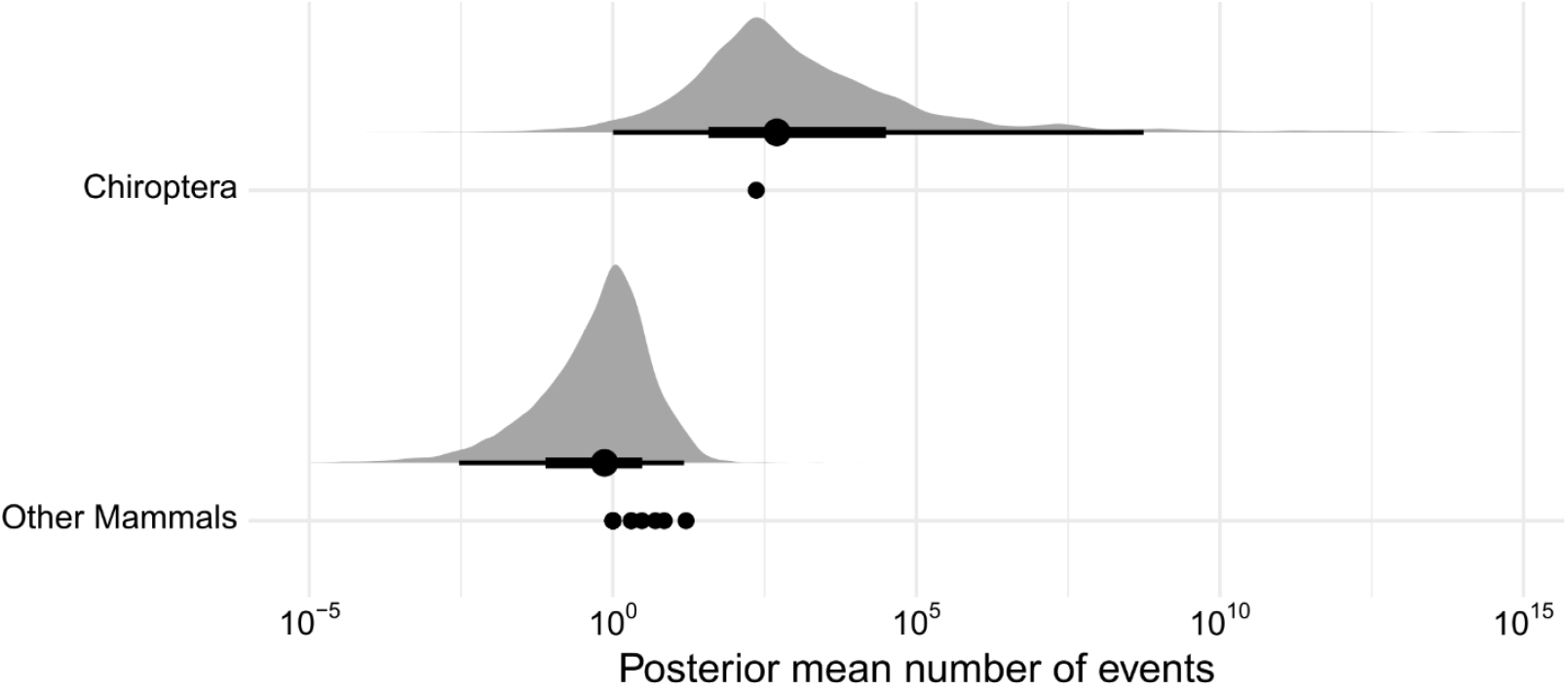
Posterior distributions of group category (bats vs. non-bat eutherian mammals) on horizontal TE transfer counts. A constant of 1 was added to HTT counts for plotting to show the wide range of posterior estimates, which spans many orders of magnitude. For each coefficient: black dots show median, thin lines show the 95% posterior probability, thick lines show the 66% posterior probability, and gray shows the posterior density of the estimates. Black dots show the observations on which the models were based.

### Varying HT Patterns and Rates in Chiroptera

We explored large scale patterns of HT within bats by mapping the 229 putative HT events onto a bat phylogeny based on the presence/absence patterns of each element and its estimated average age (Fig. 4, Table S6). As expected, there were far more putative HT events in yangochiropterans than in yinpterochiropteran lineages (170 and 59, respectively) but the distribution is exceptionally uneven within each clade. More than a third of all HT in Yinpterochiroptera are unique to *Craseonycteris thonglongyai*, with only two relatively ancient examples occurring in Pteropodidae. Similarly, within Yangochiroptera a large majority (n = 134, 78.8%) of HT events involve only vespertilionid bats. Interestingly, 8 different elements appear to have independently invaded both Vespertilionidae and either *C. thonglongyai* or the Rhinolophoidea ancestral branch (Table S5, S6), though it is unclear if these represent initial HT into one bat clade followed by HT between bats, a pair of independent HT from outside Chiroptera into different bat clades, or some combination thereof. Searches for orthologous insertions of the eight HTTs among representative species (*Hipposideros galeritus, C. thonglongyai, Myotis myotis*, and *Pipistrellus pipistrellus*), yielded zero matching orthologous insertions.

**Fig. 4.**
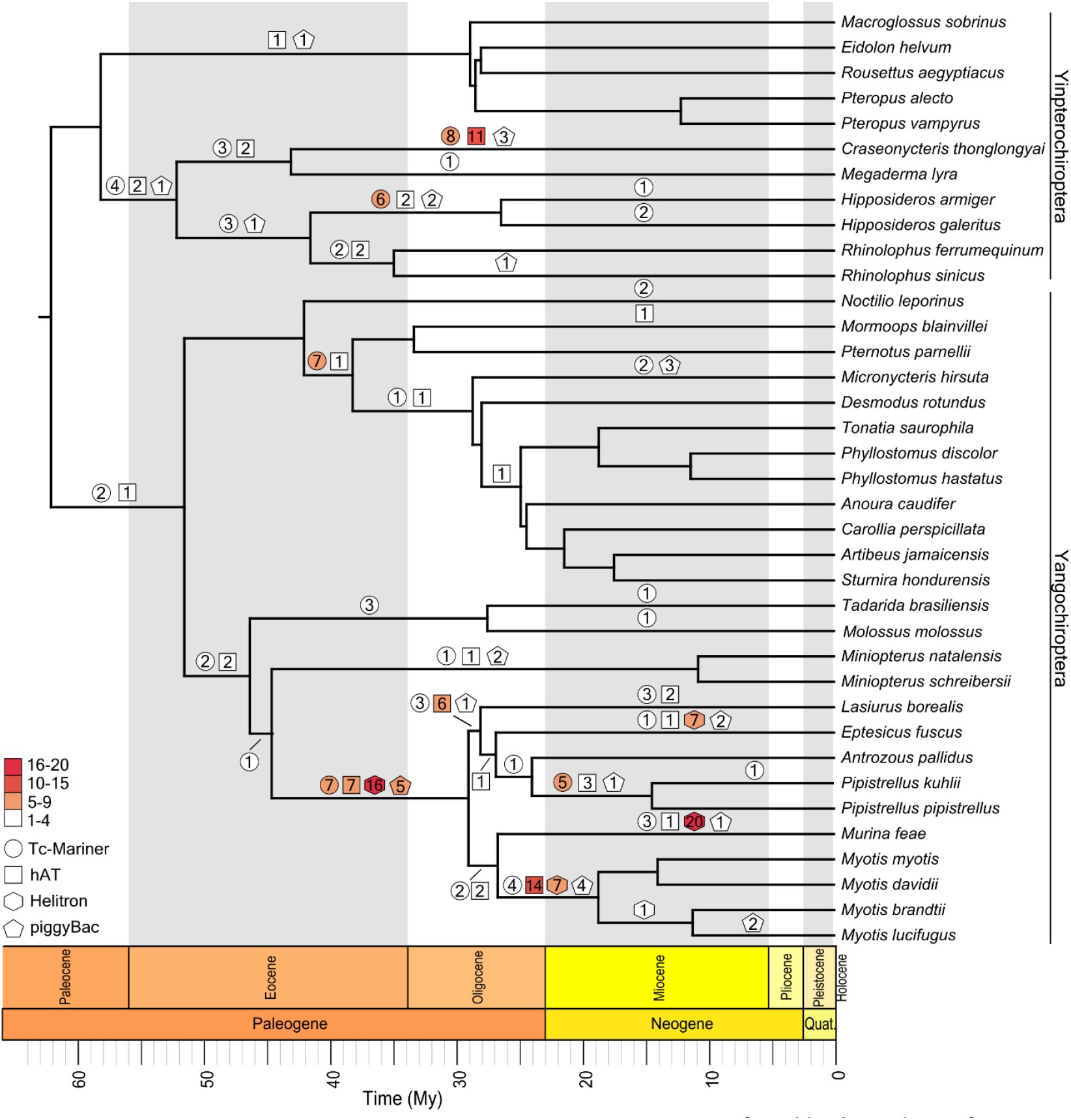
Horizontal transfer of DNA transposons within Chiroptera. Inferred horizontal transfer (HT) events of 221 unique transposable elements from Tc-Mariner (circle), hAT (square), Helitron (hexagon), or piggyBac (pentagon) families are labelled on corresponding branches. Shape color indicates numerical range of putative HT events on a given branch: white = 1-4 elements, pink = 5-9, red = 10-15, dark red = 16-20; number of events included within each marker. Phylogeny is scaled by estimated divergence times in millions of years (My). HT event branch assignment inferred from presence/absence patterns and the element’s average age. Phylogenetic relationships are based on Foley et al. (2022) and Amador et al. (2018); estimated lineage divergence times (Table S10) taken from TimeTree (Kumar, et al. 2022).

We then calculated HT event rates for bat lineages. Yangochiropterans had almost double the average HT rate of yinpterochiropterans, with a rate of 0.277 versus 0.146 putative HT/My, respectively (Table S9). However, we found a broad range of HT rates within both groups. Within Yinpterochiroptera, rates ranged from 0.023 for *Megaderma lyra* to 0.512 for *C. thonglongyai*. The ancestral branch for Hipposideridae and Rhinolophidae had the second-highest rate at 0.244. Within Yangochiroptera, rates varied between 0.022 at the ancestral branch for Miniopteridae and Vespertilionidae and 1.593 at the ancestral branch for the four *Myotis* species, which was also the highest HT rate within examined lineages. The second-highest rate within Yangochiroptera was in the ancestral Vespertilionidae branch, at 1.215 (Table S9).

Within bats, Hipposideridae, Rhinolophidae, and Vespertilionidae are among the most species rich clades while also exhibiting some of the highest TE diversity. This raises the question of a relationship between species richness and HT events. The relationship between species diversity and HT events was indeed stronger than for mammals more generally (Fig. 1). However, the relationship between species richness and HT events, despite considerable variation across TE types, proved to be statistically unsupported (Fig. S4, Table S11) but intriguing. This was also the case for only young TE counts (Fig. S5, Table S12). By increasing statistical power, additional data has the potential to influence future understanding of this relationship.

## Discussion

Our results, in combination with those of Christmas et al. (forthcoming), indicate that bats are a hotspot for horizontal transfer of DNA transposons within mammals. This was a broad-scale, computational approach to identify HTT and we used several conservative search thresholds that excluded candidate HT DNA transposons with low copy number (<90 annotated insertions) in bats, such as *Helibat1* and *SPIN_Ml*, both previously identified as HTT with limited distributions (Pace, et al. 2008; Thomas, et al. 2011). We also excluded many highly similar elements to avoid inflation from vertically diversifying elements, including highly similar deletion products. This could have yielded false negatives in both our mammalian targets and other eukaryotes. Further research into potential vectors such as eukaryotic parasites and viruses will require less conservative methods to detect low copy or fragmented elements. Despite these limitations, we found several hundred HT events, which likely are an underrepresentation of the number of HT events that have occurred within Chiroptera, particularly as HT is more likely than vertical persistence of DNA transposons (reviewed in Feschotte and Pritham 2007; Wells and Feschotte 2020). In comparison to other mammals, bats have far more HT events, and substantially higher recent DNA transposon accumulation, even when compared to mammals known to have experienced HTT, such as *Otolemur garnettii, Microcebus murinus*, or *Echinops telfairi* (Fig. 1, Fig. 3, Table S2, Table S8). While our searches did identify four species with higher than the mammalian average recent DNA transposon accumulation, these instances are clearly exceptions among non-bat eutherians and not the rule.

To better clarify the distributions and impacts of these HT events, more even sampling across bat lineages is required, particularly within large species complexes. For example, the genus *Rhinolophus* consists of ∼100 species divided among 15 species groups (Csorba, et al. 2003; Stoffberg, et al. 2010; Demos, et al. 2019), but was represented by only two genome assemblies. Since most genera are only represented by a single species, it should be noted that HT events mapped to terminal branches may represent HTs into a common ancestor of multiple species rather than our representative terminal species. That said, underrepresentation within genera would not explain the numerous lineage specific HTs of *C. thonglongyai* (26), which is a monotypic genus.

Consistent with the TE-Thrust hypothesis, most inferred HT events in Fig. 4 map to families or genera that have undergone rapid diversification. Owing to their potential for genomic innovation, TE expansions in a genome represent an opportunity for those genomes to gain variation that could lead to adaptive opportunities (Oliver and Greene 2011, 2012), giving rise to the TE-Thrust hypothesis. HT events are concentrated at the base of *Hipposideros* and *Rhinolophus* (Foley, et al. 2015), which have 90 and 106 recognized species, respectively, and Vespertilionidae (Lack and Van Den Bussche 2010), which currently consists of 512 species, and basal lineages within it, such as genus *Myotis*, which comprises 131 species (Simmons and Cirranello 2020, accessed 4 September 2021). Thus, intermittent HT and subsequent bursts of TE amplification correspond to diversification of several large clades across Chiroptera. The TE-thrust hypothesis also proposes a ‘Goldilocks Zone’ of TEs and evolutionary potential: too little TE activity results in evolutionary stasis, too much would cause detrimental genomic instability, but moderate amounts of TE activity and accumulation can allow genomic dynamism and potentially rapid lineage evolution and diversification (Oliver and Greene 2011, 2012). The data we present is consistent with these predictions. Some bat lineages, having experienced an influx of highly successful DNA transposons, may have exploited the increased genomic diversity to aid their expansion into multiple niches. Alternatively, higher species richness could lead to more HT events due to increased ecological interactions with potential HT sources and/or vectors, which could synergize with initial HT-driven diversification. Or environmental heterogeneity may promote speciation and HT, without HT directly impacting species diversification. This seems less likely given documented Helitron capture of host promoters and exons in *Myotis* (Thomas, et al. 2014). Helitron-driven tissue-specific nuclear gene transcription was shown in *Myotis brandtii* (Grabundzija, et al. 2016), and Cosby et al. (2021) identified numerous DNA transposase-gene fusions with broad gene regulatory functions that vary across bat clades, including two fusion genes specific to vespertilionids. However, we did not find statistical support for associations between horizontally transferred elements and descendent species richness, or young (≤50 My) TE accumulation and species richness, likely due to the few bat species sampled and the high variance of species richness represented by each of our focal taxa. We plan to address this in the future as additional high-quality genome assemblies are released and statistical power is increased.

While we do not know why bats are hotspots for HT, HT-associated TE diversity and accumulation, our results may indicate a higher tolerance for TE activity in bats. Possible factors influencing this presumed tolerance could include adaptations in DNA repair pathways and expression (Seim, et al. 2013; Zhang, et al. 2013; Foley, et al. 2018; Huang, et al. 2019) allowing higher TE loads. Tolerance may also have been influenced by the potential adaptations in bat immune responses that allow them to experience low viral loads but many circulating viruses with little apparent negative effects and rapid viral spreading in hosts (Subudhi, et al. 2019; Brook, et al. 2020; Jebb, et al. 2020; Irving, et al. 2021; Moreno Santillán, et al. 2021). As viruses are likely candidates for transferring TEs (Gilbert, et al. 2010; Thomas, et al. 2010; Gilbert, et al. 2014; Gilbert, et al. 2016; Gilbert and Feschotte 2018), variability within and across bat lineages in these immune-related gene expansions and losses (Moreno Santillán, et al. 2021), diversity of viruses present (Jebb, et al. 2020), as well as impacts of variable geographic proximity (Peccoud, et al. 2017) may help explain the higher frequency of HTT in chiropterans and variability of HT success across bat lineages.

Differential bat ecology may also represent part of the answer. Previous studies have implicated blood feeding arthropods such as *Rhodnius prolixus*, an insect vector of Chagas disease, as a vector for HT (Gilbert, et al. 2010; Matthews, et al. 2011). Herbivorous bats have significantly less recent DNA transposon accumulation than carnivorous species (Osmanski, et al. 2022). These observations suggest insectivorous species may be more susceptible to HT than species with other dietary habits. And indeed, the clade of bats exhibiting the highest rate of putative HT in our study is the family Vespertilionidae, which is almost exclusively insectivorous (Nowak 1999; Fenton and Bogdanowicz 2002; Morales, et al. 2019). *C. thonglongyai*, rhinolophids, and hipposiderids are also insectivorous (Arbour, et al. 2019; Pavey 2021) and stand out as exceptional genomic habitats for HT of DNA transposons. Yet despite their openness to HT, only a handful of types have been successful and with the emphatic exception of Helitrons in vespertilionids, bats do not seem to have much more diversity in DNA transposons compared to other eutherians. Why this is the case is still unclear.

The potential impacts of these HTT on bat genome evolution cannot be understated. TEs generally are a potent source of genomic variation that can impact genes and genome structure in numerous ways (Schaack, et al. 2010; Oliver and Greene 2012; Casacuberta and González 2013; Gilbert and Feschotte 2018). Studies in other mammals have shown low conservation of regulatory sites, and TEs play critical roles in restructuring regulatory networks by contributing lineage-specific transcription factor binding sites and regulatory elements (Wang, et al. 2007; Kunarso, et al. 2010; Schmidt, et al. 2012; Chuong, et al. 2013; Jacques, et al. 2013; Sundaram, et al. 2014; Notwell, et al. 2015; Trizzino, et al. 2017; Judd, et al. 2021). DNA transposons are no exception. Previous work has shown Helitron-mediated exon and promoter shuffling and substantial genome inflation within bats (Thomas, et al. 2014), as well as transposon cooption events resulting in gene fusion and changes in gene network regulation (Cosby, et al. 2021). DNA transposons are well suited to exaptation into transcription factors, as their encoded transposase proteins, a DNA binding domain and a catalytic nuclease domain, can be domesticated or repurposed for host cellular functions (Feschotte and Pritham 2007). Known host-transposase fusion genes include *GTF2IRD2* in placental mammals (Tipney, et al. 2004), *SETMAR* and *CSB-PGBD3* in primates (Cordaux, et al. 2006; Newman, et al. 2008), and *KRABINER* in Vespertilionid bats (Cosby, et al. 2021).

We note that a weakness of our study is the identification of only a few potential donor/recipient relationships to the species level. This, however, is to be expected given the paucity of animal genome assemblies available to search. Only several thousand animal genomes are available of the ∼7.8 million animal species currently estimated to exist (Mora, et al. 2011). Thus, while determining the likely HT partner in any given HT event would be ideal, doing so in all cases is difficult. We point out that, given our current understanding of evolutionary processes, the sudden appearance of multiple intact sequences with the hallmarks of DNA transposons in a lineage is likely the result of HT.

The observations presented here suggest that HTT events involving Class II transposable elements contribute to bat genomic diversity to a degree not found in other mammals. The cause of this propensity toward DNA transposon invasion is currently a mystery but future investigations may reveal the genomic characteristics that make one species more or less likely to be a safe harbor for horizontally transferred TEs. Regardless of the reasons and mechanisms behind the multiple invasions, the correspondence between high rates of HTT events and species radiations in several large bat clades suggests that HTT activity facilitates genomic innovation and taxonomic diversity. Our results shed new light on the extent of HTT in bats, but not the impacts of each example or lineage. More research is needed to clarify the specific roles that these TE expansions have played in bat diversification and genome evolution.

## Materials and Methods

### 1.1 Taxon Selection

We examined 37 bat genome assemblies and 214 other eutherian mammal assemblies for this work (Table S1). These included assemblies from the Zoonomia sequencing effort (Genereux, et al. 2020), publically available assemblies, and from other sources such as the Bat1k consortium (Jebb, et al. 2019; Wang, et al. 2020; Moreno Santillán, et al. 2021). In cases where species were represented by individuals in the Zoonomia project, but the assemblies generated by other efforts were of higher quality, we replaced the Zoonomia assemblies with the alternates (Table S13). We used a combination of PacBio, Bionano, HiC, and Illumina sequencing to generate high quality assemblies for *Eptesicus fuscus* and *Antrozous pallidus* (see Supplemental Methods).

### 1.2 Annotation of Mammalian Transposable Element Insertions

We used the curated *de novo* transposable element (TE) consensus sequence library described in Osmanski et al. (2022) to annotate TE insertions in all selected species using RepeatMasker v4.1.2-p1 (Smit, et al. 2013-2015) with the RMBlast search engine. Output was processed using RM2Bed.py, a utility in the RepeatMasker package, with TE insertion overlap resolution by lower divergence values (-o lower_div). TE insertion accumulation and temporal distributions were visualized using matplotlib (Hunter 2007) in Python v3.7.6. We estimated individual TE insertion ages by calculating species-specific neutral mutation rates for all lineages within the last ∼50 My using pairwise branch lengths from Foley et al. (2022) and median divergence times for each species versus an outgroup mammal taxon from TimeTree (Kumar, et al. 2022). We then evaluated the TE content of the 213 non-bat eutherian mammals and selected the four species with the highest recent DNA transposon accumulation to compare to bats, as well as four other species representing eutherian orders closely related to Chiroptera. Annotations for rolling-circle elements (Helitrons) in bat species outside of Vespertilionidae were excluded from these visualizations, as these are known to be false positives, as discussed in Osmanski et al. (2022).

### 2.1 Identification of Putative Horizontally Transferred Class II TEs Involving Chiroptera and Other Mammals

We selected DNA/RC elements with ≥90 annotated copies in at least one bat species as our initial set of HT candidates. We then used the library consensus sequences (107) of this initial TE set as queries in BLAST searches utilizing what we refer to as the 90-90-90 rule (described below), a more conservative version of the 80-80-80 rule developed by Wicker et al. (Wicker, et al. 2007). We searched for TE copies meeting our conservative criteria of present in the genome assemblies of one or more bat species. To identify any additional eukaryote involvement, we performed BLAST searches of these elements across all available eukaryote genome assemblies in the NCBI databases.

Putative horizontally transferred transposable elements (HTT) were defined as TE insertions annotated in an assembly with <90 insertions called in closely related species. We narrowed our search for HTT to DNA transposon and rolling-circle transposons with ≥90 copies annotated by RepeatMasker in one or more bat species. We then used the same TE consensus sequences as queries for blastn searches (BLAST+ v2.11.0 (Camacho, et al. 2009)) in said bat genomes and implemented the 90-90-90 rule to identify potential HTTs. The criteria of the 90-90-90 rule are 1) the element must be ≥90 bp in length, 2) share ≥90% sequence identity with one another, and 3) have a total ungapped length matching ≥90% of the consensus sequence. To further exclude potentially erroneous hits from similar elements harboring short insertions, the element copies must have been ≤10% longer than the query consensus sequence length. We also excluded potential duplicate elements or vertically diversifying elements with ≤5% sequence divergence using the cross_match utility of Phrap v0.990319 (Gordon 2003). Similarly, to account for and exclude DNA transposon deletion products, we used the same query consensus sequences as before to perform a modified CD-HIT (Storer, Hubley, Rosen and Smit 2021) search for candidate HTT sequences that cluster together. This search performs two successive cd-hit searches. The first clusters elements ≥ 90% identical, the second search adds elements >80% similar to existing clusters or generates new ones. Elements that clustered together and had overlapping presence/absence patterns across bat species were collapsed into a single presumed HT event.

We then performed a final manual curation by comparing alignments of candidate HTT consensus sequences to all other elements in the TE consensus library from section 1.2 to identify any deletion products that were not identified in the previous clustering step. To estimate the age of each TE insertion within a species, we calculated modified Kimura two-parameter (K2P) distances for each TE copy compared to the library consensus sequence using RepeatMasker’s alignAndCallConsensus and Linup utilities (Smit, et al. 2013-2015). We then mapped the HT events onto a phylogenetic tree of our 37 bat species based on the presence/absence pattern of the putative HT elements from our filtered blastn results and their average ages. TE ages were calculated per species using the average K2P distance and the species-specific neutral mutation rates. The phylogenetic tree was built based on Foley et al. (2022) and Amador et al. (2018), and used a combination of non-conflicting average or median divergence estimates from TimeTree (Table S10) (Kumar, et al. 2022), accessed 3 September 2021.

### 2.2 Orthologous TE Insertion Searches within Mammalia

To identify possible orthologous copies of putative HTTs, we performed pairwise orthologous site searches between twenty-eight bats species and two mammal outgroups, *Bos taurus* and *Equus caballus*, using Zoonomia’s 241 mammal genome alignment (Genereux, et al. 2020). With the exception of *Noctilio leporinus*, the other eight bat species not present in the genome alignment were represented by other members in the same family, if not the same genus. For each of the twenty-eight bat species, we generated a BED file of the coordinates of each copy of a putative HTT in the final dataset from 2.1 with 50 bp flanking sequence on either end. We then identified the orthologous sections of the outgroup genomes with the utility halLiftover, and merged all close (≤2 bp) coordinate hits for the same TE copy into a single hit using BEDTools sort and mergeBed (Quinlan and Hall 2010). We then performed a series of TE annotations for all orthologous sites in the target outgroup species, first using RepeatMasker (Smit, et al. 2013-2015) with a combined mammalian TE consensus library of ancestral mammal repeats from the Dfam database v.3.6 (Storer, Hubley, Rosen, Wheeler, et al. 2021) and our original library from section 1.2. Any annotations matching one of the 221 putative HTTs were then subjected to an additional annotation and alignment with the cross_match utility (Gordon 2003). Any cross_match annotations matching one of the 221 putative HTTs were then manually checked for 1) TE identity match to the copy at the bat site, 2) alignment size and score, and 3) site alignment to bat species (e.g. were there large (>1000 bp) gaps). The same process of pairwise orthologous site searches was performed with representative species for mammal groups harboring any of putative HTTs, which included *Microgale talazaci* (Afrosoricida), *Tupaia chinensis* (Scandentia), *Nycticebus coucang* (Lemuriformes), *Crocidura indochinensis* (Eulipotyphla). These mammals were paired with representative bat species: *Hipposideros galeritus, Myotis myotis, Murina feae*, and/or *Pipistrellus pipistrellus*. We also performed orthologous site searches between representatives of the two bat suborders: Yinpterochiroptera (*Craseonycteris thonglongyai, Hipposideros galeritus*) and Yangochiroptera (*Myotis myotis, Pipistrellus pipistrellus*).

### 2.3 Identification of Putative Horizontally Transferred TEs Outside of Mammalia

After identifying HT events, we applied the above methodology to identify possible HT events between Chiroptera and non-mammal eukaryotes. We performed blastn searches of the eukaryotic reference genome database (accessed 6 April 2021 (Camacho, et al. 2009)), excluding mammals, using the consensus sequences from the putative chiropteran HTTs as our query input. To reduce false negatives in distantly related taxa, we used the criterion of ≥90 full-length or near full-length copies for non-autonomous elements, and a lower threshold of ≥50 copies for autonomous elements. As non-autonomous copies tend to make up the majority of DNA transposon insertions (Lohe, et al. 1995; Feschotte and Pritham 2007; Muñoz-López and García-Pérez 2010), this threshold is more likely to detect true evolutionarily recent HTT in more distantly related organisms. To identify autonomous elements, we searched for open reading frames (ORFs) via the getorf utility of EMBOSS v6.6.0 (Rice, et al. 2000) in species-specific consensus sequences of the putative HTT generated from a custom script, extend_align.sh, which is available on Github (https://github.com/davidaray/bioinfo_tools). We identified transposase ORFs by performing blastx searches.

### 2.3 Testing for Associations with Species Richness

Two sets of analyses were conducted. First, we tested the association between horizontally transferred TEs and fraction species richness modelling both these variables with errors. Then, we modelled fraction species richness as a function of cumulative young (≤50 My) TEs (see Supplemental Methods for details).

## Supporting information

https://david-ray-7mp3.squarespace.com/s/Paulat_et_al_Bat_HT_supplementary_files.zip

## Acknowledgements

This project was supported by the National Science Foundation (grant numbers DEB 1838283 and IOS 2032006 to D.D.M.S. and D.A.R., DEB 1838273 and DGE 1633299 to L.M.D.); and National Institutes of Health (grant numbers R01HG002939 and U24HG010136 to J.M.S., R.H., A.S., Jeb R.), NHGRI R01HG008742 to Z.C.); and the Irish Research Council (grant number IRCLA/2017/58 to E.C.T.); and the Science Foundation Ireland (grant number 19/FFP/6790 to E.C.T.); and Max Planck Research Group awarded by the Max Planck Gesellschaft to S.C.V.; and Human Frontiers Science Program (grant number RGP0058/2016 to S.C.V.); and UK Research and Innovation (grant number MR/T021985/1 to S.C.V.); and the Swedish Research Council Distinguished Professor Award to K.L.T. The High-Performance Computing Center at Texas Tech University and the SeaWulf computing system at Stony Brook University provided computational infrastructure throughout the work.

## Author contributions

N.S.P. and D.A.R. contributed to conceptualization, design, data analysis and interpretation, and drafted the manuscript. N.S.P., J.M.S., A.B.O., K.A.M.S., J.K., J.R.G., M.H., C.G., C.C., J.R., JebR., R.H., A.S., and D.A.R. participated in library validation and curation. Genome assembly was accomplished by M.P., T. B., and M.H. Methods and interpretation were contributed by N.S.P., J.M.S., A.B.O., D.A.R., L.M.D., D.R., K.L-T., E.K.K. and D.D.M.S. All authors contributed to review and editing of the final manuscript. All authors gave final approval and agreed to be accountable for all aspects of the work.

## Data Availability Statement

All assemblies are available on Genbank (see ST1 for accession numbers). TE consensus sequences, and their seed alignments, are available via the Dfam database. All other data is available in the supplementary materials; code used in the analysis is available at: github.com/daray/bat_ht.

