## Supplementary material for "Chiropterans are a hotspot for horizontal transfer of DNA transposons in Mammalia": https://david-ray-7mp3.squarespace.com/s/Paulat_et_al_Bat_HT_supplementary_files.zip: Paulat_et_al_Bat_HT_supplementary_text_MBE_march02.docx

**Supplementary Materials**

**Supplemental Methods**

***De Novo Genome Assembly***

Contig Assembly

For both *Antrozous pallidus* and *Eptesicus fuscus*, initial contigs were assembled from PacBio subreads using DAmar (<https://github.com/MartinPippel/DAmar>, commit: 3624a8c) and two rounds of polishing with gcpp v2.0.2 (https://github.com/PacificBiosciences/gcpp). Remaining haplotypic duplications in the primary contig set were then removed using purge-dups v.1.2.3 (Guan, et al. 2020).

*Antrozous pallidus* Scaffolding

Initial scaffolding for *A. pallidus* was performed using 10X Genomics linked-reads. First, reads were mapped to the contigs using Longranger v2.2.2 (Marks, et al. 2019) and scaffolded using Scaff10X v4.2 (<https://github.com/wtsi-hpag/Scaff10X>) . Next, we used optical maps from Bionano DLE1-labelled DNA molecules. A Bionano assembly was first produced from optical-mapped reads using Bionano Solve assembly v3.5.1 with arguments non-haplotype and no-Extend-and-Split. The initial scaffolds from Scaff10X were then further scaffolded using Bionano Solve Hybrid Scaffold v3.5.1. Finally, HiC reads were mapped to the scaffolds produced by Hybrid Scaffold using bwa-mem v0.7.17 (Li and Durbin 2009, 2010). We then followed the VGP scaffolding pipeline with salsa2 v2.2 (Ghurye, et al. 2019; Rhie, et al. 2021). Finally, we manually curated the scaffolds to join those contigs missed by salsa2 and break those joins which were spuriously created.

After initial scaffolding, we closed assembly gaps using the PacBio CLR data. To this end, we mapped the original subreads.bam files to the scaffolded assembly using pbmm2 v1.3.0 with arguments --preset SUBREAD -N 1. Based on the read-piles created by reads spanning across gap regions, we can create a consensus sequence to replace the N sequences in our genome. We used gcpp v2.0.2 to polish gap regions and their 2 kb up/downstream flanks. We then replaced the assembly gap and flanking region with those regions that were polished with high-confidence by arrow (no N's remaining in the polished sequence and no lower-case a/c/g/t).

*Eptesicus fuscus* Scaffolding

HiC reads were mapped to the primary contigs using bwa-mem v0.7.17 and the resulting alignments were further processed with pairtools v0.3.0 (https://github.com/open2c/pairtools) according to the Omni-C filtering pipeline (<https://omni-c.readthedocs.io/en/latest/fastq_to_bam.html>). The resulting deduplicated bam file sorted by read names and YaHS v1.1 (Zhou, et al. 2022) was used to scaffold the contigs. Finally, we manually curated the scaffolds to join those contigs missed by YaHS and break those joins which were spuriously created.

Assembly Polishing

To create a higher-accuracy genome at the level base accuracy, we polished the resulting assemblies using the 10X linked-reads (*A. pallidus*) and Illumina reads (*E. fuscus*) as in the VGP pipeline (<https://github.com/VGP/vgp-assembly/tree/master/pipeline/freebayes-polish>) (Rhie, et al. 2021). First 10X reads were mapped to the *A. pallidus* genome using Longranger v2.2.2, while Illumina reads were mapped to the *E. fuscus* genome with bwa-mem v0.7.17. In both cases, variants were called using Freebayes v1.3.2 with argument -g 600 to ignore regions with coverage over 600X. Next variants were filtered using bcftools v1.12-21 for variants with quality score greater than 1 and genotype of homozygous alt (AA) or heterozygous (Aa): bcftools view -i 'QUAL>1 && (GT="AA" || GT="Aa")'. A consensus was then called using bcftools consensus, taking the longest allele in heterozyous cases: bcftools consensus -i'QUAL>1 && (GT="AA" || GT="Aa")' –Hla. This was performed twice; the *A. pallidus* consensus in the first round changing 3,766,332 bases and in the second 217,026 bases.

For *A. pallidus*, we estimated the QV of this assembly after two rounds of polishing to be 33.6 using merqury v1.0 (Rhie, et al. 2020). Finally, the curated chromosomes were phased by applying an adapted version of the DipAsm pipeline (Garg, et al. 2021).

For *E. fuscus*, to further increase the scaffold accuracy, we applied merfin v1.0 (Formenti, et al. 2022). Using merqury v1.0, we estimated the QV of this assembly after two rounds of polishing to be 38.0.

***Species Richness Association Testing***

The first set of analyses comprised three steps. Initially, we estimated the species richness represented by each branch in the TE phylogeny, then analyzed the association between TE counts and richness while accounting for errors in both variables and the phylogenetic structure of errors.

We used a phylogenetic approach to determine the species richness represented by each branch. We first used the most comprehensive species-level phylogeny of bats by Shi and Rabosky (Shi and Rabosky 2015), which includes most of the species in the sample or close relative of those when not sampled. While we could assign the family-level values to each species, this approach would flatten the variation across species found in TEs and the representativeness of the branches. Instead, we partitioned the species-level tree into subtrees then evaluated which subtrees had the target species. The species richness was estimated by counting tips in the largest subtree representing one and only one of the species in the TE dataset. Then, the proportion of species represented was calculated by dividing the number of leaves by the sum for all species in the sample.

To estimate the association between species diversity and TE variables, we generated linear models. However, observations including multiple species are not independent (Felsenstein 1985), and therefore cannot be analyzed using standard statistical methods all of which assume independence among observations. Therefore, we adopted methods that enable accounting for the non-independence among errors in the linear models by modeling this structure based on phylogenetic distances among all species-level observations. As with standard linear models, these methods estimate the relationship between variables as coefficients that multiply quantities (e.g., TE counts) or categories (e.g., bat observations, vs. all others). The sign (positive or negative) of the coefficients indicates the quantitative relationship between variables, with coefficients of 0 consistent with no association of one variable on the other.

A hierarchical Bayesian approach was adopted to estimate the species-specific structure of errors while estimating error for both the Poisson-distributed TE counts and the beta-distributed proportion of richness represented by each branch. A hierarchical approach is often called a mixed model in the literature, with cluster-specific effects called “random”, and sample-wide effects called “fixed”. As different fields apply random and fixed to different levels of the hierarchy, here we adopt the language of cluster-specific and sample-wide effects (Gelman 2005). Analyses begin by modelling the proportion of richness as a beta-distributed variable (Douma and Weedon 2019):

$$y_{1i}\sim Beta(\mu,\phi)$$

In which $\mu$ is the mean, and $\phi$ relates to the variance such that:

$$var\left[ y_{1} \right]=\frac{\mu(1-\mu)}{1+\phi}$$

Given observations *Y_1_*, and covariate *Y_2_*:

$$logit\left( \mu\right)=\log\left( \frac{\mu}{1-\mu} \right)={\beta Y}_{2}$$

Instead of a typical regression, in which the covariate is modeled without error, our analyses accounted for the error in TE counts by modeling the latter as a negative binomial-distributed variable:

$$y_{2i}\sim negative binomial(\lambda=\exp\left( l_{i} \right), {pr}_{m,t})$$

In which $\lambda$ is the rate or mean of the Poisson distribution, exp is the n verse link function that enables the inclusion of phylogenetic errors (Hadfield and Nakagawa 2010), and $\frac{1-pr}{pr}$ defines a rate parameter for the gamma distribution that defines the mixture of Poisson distributions, which relaxes the expectation of equality of mean and variance of the Poisson distribution. As a result, the negative binomial distribution is usually a better fit to observational data (O'Hara and Kotze 2010). With a linear model applied to *l*:

$$l=\beta_{0}\theta+e$$

In which $\beta_{0}$ represents the intercept, as there were no predictors included, $\theta$ represents both species specific and sample-wide effects, and e is a vector of residuals. The sample-wide coefficient is normally distributed and given by:

$$\beta_{0}\sim N(0,\sigma_{B}^{2}I)$$

In which the variance $\sigma_{B}^{2}$ is large, reflecting diffuse prior knowledge. The phylogeny-based species-specific effects that account for relatedness are also normally distributed and given by:

$$a\sim N(0,\sigma_{a}^{2}A)$$

Which replaces the identity matrix *I* with the phylogenetic relationship matrix *A*. With no other levels, the is modeled by $\sigma_{e}^{2}$ in:

$e\sim N(0,\sigma_{e}^{2}I)$.

A similar approach was used to fit the phylogenetic structure of errors to the beta regression. In contrast with horizontally transferred TE counts, cumulative TE counts generally spanned a couple of orders of magnitude. To avoid overfitting these analyses, we modeled errors only for the beta-distributed proportions of richness. To span the order-of-magnitude variation in the TE accumulation counts, this predictor was log10 transformed and then scaled. Similarly, we fitted models of HTT events (a negative-binomial-distributed variable) across 19 mammalian orders (Table S8) to compare the single bat observation to the HTT counts from all other mammals.

To implement Bayesian sampling for these analyses, we used brms (Bürkner 2017), a package that enables coding models in R for implementation in the stan statistical language (Carpenter, et al. 2017). We ran separate multivariate models for each of the TE counts, with the proportion of richness as a function of the TE count and the count itself as a response. The covariance matrix A was obtained from the variance covariance matrix of the dated phylogeny of sampled species from 2.1. Models ran four separate chains using a Hamiltonian Monte Carlo approach. Compared to other Bayesian implementations, the HMC approach saves time in sampling parameter spaces by generating efficient transitions spanning the posterior based on derivatives of the density function of the model.

**Supplemental Results**

***TE Accumulation Outliers***

*C. thonglongyai* and *Me. lyra* are outliers (highest 2.5% of data) with the highest total DNA transposon accumulation (8.4% and 6.3% of their genome, respectively). *C. thonglongyai* and *Pipistrellus pipistrellus* are also outliers for highest accumulation in the last 50 My, with young DNA transposons comprising 0.34% and 0.24% of their genomes, respectively. Outgroup mammals make up the low-end outliers (lowest 2.5% of data) for both total and younger DNA transposon accumulation (*Erinaceus europaeus*: 1%, *Uropsilus gracilus*: 2.2%, and *Equus caballus*: 0.0039%, *Diceros bicornis*: 0.0026%, respectively). For RC elements, *Murina feae* is an outlier for high total and younger RC accumulation, constituting 19.7% and 10.3% of its genome, respectively. Other vespertilionid bats have the next highest RC genome content; for total RC content, *Antrozous pallidus* with 14.7%, and for younger RC content, *Eptesicus fuscus* with 5.1%. No RC elements have invaded other mammalian orders.

***Effects of Genome Assembly Quality on TE Annotation***

We should also note that although some differences in TE accumulation among species may be an artifact of varying genome assembly qualities, these do not explain our results. Across all of the mammal assemblies there was no clear or consistent trend of TE proportions based on assembly N50 or BUSCO scores, though outliers for high total genomic TE content tended to be assemblies with low N50 or low BUSCO scores (Osmanski, et al. forthcoming). We observed only minimal differences in TE proportions between closely related species. For example, the four representatives of the genus *Myotis* diverged between ~20 and 10 Mya (Stadelmann, et al. 2007; Lack, et al. 2010; Ruedi, et al. 2013), and *Myotis myotis* has a higher quality assembly (N50 = 94.4 Mb, BUSCO = 97.9%) than the other three *Myotis* species (mean N50 = 3.7 Mb, mean BUSCO = 92.5%). While there was some minor variation in TE content, major trends and temporal patterns are consistent, and *Myotis myotis* actually had the highest TE content of the four species. While genome fragmentation might have affected observed TE proportions, this likely would have only led to underrepresentation or false negatives for our HTT analyses, since we used only insertions ≥90% consensus sequence length.

***No Significant Associations between Putative HT DNA Transposons and Species Richness***

Models ran for at least 10,000 generations, with at least 20% of the generations sampled as burn in. All models ran until estimated sampling sizes for posteriors exceeded 1000, the potential scale reduction factor was no greater than 1.05 (indicating convergence across chains), and there were no divergent transitions after burn-in. The absence of divergent transitions indicates the sampling was unbiased. We found no significant associations between putative HTTs and species richness (fig. S4, Table S11).

***Limited Association between Young DNA Transposon Accumulation and Species Richness***

We found only a weak association with cumulative LTRs and species richness; all other categories had no significant associations (fig. S5, Table S12). This is likely due to the small sample size of bat species; more data would probably yield more meaningful results.

***Zoonomia Consortium Author List***

Gregory Andrews^1^, Joel C. Armstrong^2^, Matteo Bianchi^3^, Bruce W. Birren^4^, Kevin R. Bredemeyer^5^, Ana M. Breit^6^, Matthew J. Christmas^3^, Hiram Clawson^2^, Joana Damas^7^, Federica Di Palma^8,9^, Mark Diekhans^2^, Michael X. Dong^3^, Eduardo Eizirik^10^, Kaili Fan^1^, Cornelia Fanter^11^, Nicole M. Foley^5^, Karin Forsberg-Nilsson^12,13^, Carlos J. Garcia^14^, John Gatesy^15^, Steven Gazal^16^, Diane P. Genereux^4^, Linda Goodman^17^, Jenna Grimshaw^14^, Michaela K. Halsey^14^, Andrew J. Harris^5^, Glenn Hickey^18^, Michael Hiller^19,20,21^, Allyson G. Hindle^11^, Robert M. Hubley^22^, Graham M. Hughes^23^, Jeremy Johnson^4^, David Juan^24^, Irene M. Kaplow^25,26^, Elinor K. Karlsson^1,4,27^, Kathleen C. Keough^17,28,29^, Bogdan Kirilenko^19,20,21^, Klaus-Peter Koepfli^30,31,32^, Jennifer M. Korstian^14^, Amanda Kowalczyk^25,26^, Sergey V. Kozyrev^3^, Alyssa J. Lawler^4,26,33^, Colleen Lawless^23^, Thomas Lehmann^34^, Danielle L. Levesque^6^, Harris A. Lewin^7,35,36^, Xue Li^1,4,37^, Abigail Lind^28,29^, Kerstin Lindblad-Toh^3,4^, Ava Mackay-Smith^38^, Voichita D. Marinescu^3^, Tomas Marques-Bonet^39,40,41,42^, Victor C. Mason^43^, Jennifer R. S. Meadows^3^, Wynn K. Meyer^44^, Jill E. Moore^1^, Lucas R. Moreira^1,4^, Diana D. Moreno-Santillan^14^, Kathleen M. Morrill^1,4,37^, Gerard Muntané^24^, William J. Murphy^5^, Arcadi Navarro^39,41,45,46^, Martin Nweeia^47,48,49,50^, Sylvia Ortmann^51^, Austin Osmanski^14^, Benedict Paten^2^, Nicole S. Paulat^14^, Andreas R. Pfenning^25,26^, BaDoi N. Phan^25,26,52^, Katherine S. Pollard^28,29,53^, Henry E. Pratt^1^, David A. Ray^14^, Steven K. Reilly^38^, Jeb R. Rosen^22^, Irina Ruf^54^, Louise Ryan^23^, Oliver A. Ryder^55,56^, Pardis C. Sabeti^4,57,58^, Daniel E. Schäffer^25^, Aitor Serres^24^, Beth Shapiro^59,60^, Arian F. A. Smit^22^, Mark Springer^61^, Chaitanya Srinivasan^25^, Cynthia Steiner^55^, Jessica M. Storer^22^, Kevin A. M. Sullivan^14^, Patrick F. Sullivan^62,63^, Elisabeth Sundström^3^, Megan A. Supple^59^, Ross Swofford^4^, Joy-El Talbot^64^, Emma Teeling^23^, Jason Turner-Maier^4^, Alejandro Valenzuela^24^, Franziska Wagner^65^, Ola Wallerman^3^, Chao Wang^3^, Juehan Wang^16^, Zhiping Weng^1^, Aryn P. Wilder^55^, Morgan E. Wirthlin^25,26,66^, James R. Xue^4,57^, Xiaomeng Zhang^4,25,26^

Affiliations:
^1^Program in Bioinformatics and Integrative Biology, UMass Chan Medical School; Worcester, MA 01605, USA.
^2^Genomics Institute, University of California Santa Cruz; Santa Cruz, CA 95064, USA.
^3^Department of Medical Biochemistry and Microbiology, Science for Life Laboratory, Uppsala University; Uppsala, 751 32, Sweden.
^4^Broad Institute of MIT and Harvard; Cambridge, MA 02139, USA.
^5^Veterinary Integrative Biosciences, Texas A&M University; College Station, TX 77843, USA.
^6^School of Biology and Ecology, University of Maine; Orono, ME 04469, USA.
^7^The Genome Center, University of California Davis; Davis, CA 95616, USA.
^8^Genome British Columbia; Vancouver, BC, Canada.
^9^School of Biological Sciences, University of East Anglia; Norwich, UK.
^10^School of Health and Life Sciences, Pontifical Catholic University of Rio Grande do Sul; Porto Alegre, 90619-900, Brazil.
^11^School of Life Sciences, University of Nevada Las Vegas; Las Vegas, NV 89154, USA.
^12^Biodiscovery Institute, University of Nottingham; Nottingham, UK.
^13^Department of Immunology, Genetics and Pathology, Science for Life Laboratory, Uppsala University; Uppsala, 751 85, Sweden.
^14^Department of Biological Sciences, Texas Tech University; Lubbock, TX 79409, USA.
^15^Division of Vertebrate Zoology, American Museum of Natural History; New York, NY 10024, USA.
^16^Keck School of Medicine, University of Southern California; Los Angeles, CA 90033, USA.
^17^Fauna Bio Incorporated; Emeryville, CA 94608, USA.
^18^Baskin School of Engineering, University of California Santa Cruz; Santa Cruz, CA 95064, USA.
^19^Faculty of Biosciences, Goethe-University; 60438 Frankfurt, Germany.
^20^LOEWE Centre for Translational Biodiversity Genomics; 60325 Frankfurt, Germany.
^21^Senckenberg Research Institute; 60325 Frankfurt, Germany.
^22^Institute for Systems Biology; Seattle, WA 98109, USA.
^23^School of Biology and Environmental Science, University College Dublin; Belfield, Dublin 4, Ireland.
^24^Department of Experimental and Health Sciences, Institute of Evolutionary Biology (UPF-CSIC), Universitat Pompeu Fabra; Barcelona, 08003, Spain.
^25^Department of Computational Biology, School of Computer Science, Carnegie Mellon University; Pittsburgh, PA 15213, USA.
^26^Neuroscience Institute, Carnegie Mellon University; Pittsburgh, PA 15213, USA.
^27^Program in Molecular Medicine, UMass Chan Medical School; Worcester, MA 01605, USA.
^28^Department of Epidemiology & Biostatistics, University of California San Francisco; San Francisco, CA 94158, USA.
^29^Gladstone Institutes; San Francisco, CA 94158, USA.
^30^Center for Species Survival, Smithsonian's National Zoo and Conservation Biology Institute; Washington, DC 20008, USA.
^31^Computer Technologies Laboratory, ITMO University; St. Petersburg 197101, Russia.
^32^Smithsonian-Mason School of Conservation, George Mason University; Front Royal, VA 22630, USA.
^33^Department of Biological Sciences, Mellon College of Science, Carnegie Mellon University; Pittsburgh, PA 15213, USA.
^34^Senckenberg Research Institute and Natural History Museum Frankfurt; 60325 Frankfurt am Main, Germany.
^35^Department of Evolution and Ecology, University of California Davis; Davis, CA 95616, USA.
^36^John Muir Institute for the Environment, University of California Davis; Davis, CA 95616, USA.
^37^Morningside Graduate School of Biomedical Sciences, UMass Chan Medical School; Worcester, MA 01605, USA.
^38^Department of Genetics, Yale School of Medicine; New Haven, CT 06510, USA.
^39^Catalan Institution of Research and Advanced Studies (ICREA); Barcelona, 08010, Spain.
^40^CNAG-CRG, Centre for Genomic Regulation, Barcelona Institute of Science and Technology (BIST); Barcelona, 08036, Spain.
^41^Department of Medicine and LIfe Sciences, Institute of Evolutionary Biology (UPF-CSIC), Universitat Pompeu Fabra; Barcelona, 08003, Spain.
^42^Institut Català de Paleontologia Miquel Crusafont, Universitat Autònoma de Barcelona; 08193, Cerdanyola del Vallès, Barcelona, Spain.
^43^Institute of Cell Biology, University of Bern; 3012, Bern, Switzerland.
^44^Department of Biological Sciences, Lehigh University; Bethlehem, PA 18015, USA.
^45^BarcelonaBeta Brain Research Center, Pasqual Maragall Foundation; Barcelona, 08005, Spain.
^46^CRG, Centre for Genomic Regulation, Barcelona Institute of Science and Technology (BIST); Barcelona, 08003, Spain.
^47^Department of Comprehensive Care, School of Dental Medicine, Case Western Reserve University; Cleveland, OH 44106, USA.
^48^Department of Vertebrate Zoology, Canadian Museum of Nature; Ottawa, Ontario K2P 2R1, Canada.
^49^Department of Vertebrate Zoology, Smithsonian Institution; Washington, DC 20002, USA.
^50^Narwhal Genome Initiative, Department of Restorative Dentistry and Biomaterials Sciences, Harvard School of Dental Medicine; Boston, MA 02115, USA.
^51^Department of Evolutionary Ecology, Leibniz Institute for Zoo and Wildlife Research; 10315 Berlin, Germany.
^52^Medical Scientist Training Program, University of Pittsburgh School of Medicine; Pittsburgh, PA 15261, USA.
^53^Chan Zuckerberg Biohub; San Francisco, CA 94158, USA.
^54^Division of Messel Research and Mammalogy, Senckenberg Research Institute and Natural History Museum Frankfurt; 60325 Frankfurt am Main, Germany.
^55^Conservation Genetics, San Diego Zoo Wildlife Alliance; Escondido, CA 92027, USA.
^56^Department of Evolution, Behavior and Ecology, School of Biological Sciences, University of California San Diego; La Jolla, CA 92039, USA.
^57^Department of Organismic and Evolutionary Biology, Harvard University; Cambridge, MA 02138, USA.
^58^Howard Hughes Medical Institute; Chevy Chase, MD, USA.
^59^Department of Ecology and Evolutionary Biology, University of California Santa Cruz; Santa Cruz, CA 95064, USA.
^60^Howard Hughes Medical Institute, University of California Santa Cruz; Santa Cruz, CA 95064, USA.
^61^Department of Evolution, Ecology and Organismal Biology, University of California Riverside; Riverside, CA 92521, USA.
^62^Department of Genetics, University of North Carolina Medical School; Chapel Hill, NC 27599, USA.
^63^Department of Medical Epidemiology and Biostatistics, Karolinska Institutet; Stockholm, Sweden.
^64^Iris Data Solutions, LLC; Orono, ME 04473, USA.
^65^Museum of Zoology, Senckenberg Natural History Collections Dresden; 01109 Dresden, Germany.
^66^Allen Institute for Brain Science; Seattle, WA 98109, USA

**Supplemental Figures**

```
                 10        20        30        40        50        60        70        80        90       100
        ....|....|....|....|....|....|....|....|....|....|....|....|....|....|....|....|....|....|....|....|
PipKuh  CAGTGGTGCCTCGCATAAAGAACGCCTCGCACAGCGAACGCTGCACACAACGAACTTCATTTCATGATTCATACAACGAACTTCGTTTCACACAACGAAG
ErpCal  CAGTGGTGCCTCGCATAAAGAACGCCTCGCACAGCGAACGCTGCACACAACGAACTTCATTTCATGATTCATACAACGAACTTCGTTTCACACAACGAAG

                110       120       130       140       150       160       170       180       190       200
        ....|....|....|....|....|....|....|....|....|....|....|....|....|....|....|....|....|....|....|....|
PipKuh  TCGCCCGAGCTTCCACGACCGCTTTGCCCGAGCTTCCACGACTGCTTCGCCCGAGCTTCCACGATCGCTGCCGATGTATTGCATCCTTCCGCGCAGGCAC
ErpCal  TCGCCCGAGCTTCCACGACCGCTTTGCCCGAGCTTCCACGACTGCTTCGCCCGAGCTTCCACGATCGCTGCCGATGTATTGCATCCTTCCGCGCAGGCAC

                210       220       230       240       250       260       270       280       290       300
        ....|....|....|....|....|....|....|....|....|....|....|....|....|....|....|....|....|....|....|....|
PipKuh  TGCAGGCAGTCGTTAGTCACTGCGCTTAACTTAAGCACGTATCGCGGCAATCGTTCTTCATTACGAATTAAATCACGCACGTATCACGGCAGTCGTCCTT
ErpCal  TGCAGGCAGTCGTTAGTCACTGCGCTTAACTTAAGCACGTATCGCGGCAATCTTTCTTCATTACGAATTAAATCACGCACGTATCACGGCAGTCGTCCTT

                310       320       330       340       350       360       370       380       390       400
        ....|....|....|....|....|....|....|....|....|....|....|....|....|....|....|....|....|....|....|....|
PipKuh  TTAAGTTAAACTCAATATTTTTTTTATATCATGGCTTCTAAAAAAAGCAGGAAGGTGATTTCTGTTGAAATGAAACAGGAAATAATTAGAAGGAGTGAAT
ErpCal  TTAAGTTAAACTCAATATTTTTTTTATATCATGGCTTCTAAAAAAAGCAGGAAGGTGATTTCTGTTGAAATGAAACAGGAAATAATTAGAAGGAGTGAAT

                410       420       430       440       450       460       470       480       490       500
        ....|....|....|....|....|....|....|....|....|....|....|....|....|....|....|....|....|....|....|....|
PipKuh  GTGGGGTAAAACAGTGTGACCTCGTCAAAGAGTTTGGCCTCAGCAAGACCACCATTTTCACCATTTTGACAAATAAGGATGCAATCAAATCAGCCAAAGT
ErpCal  GTGGGGTAAAACAGTGTGACCTCGTCAAAGAGTTTGGCCTCAGCAAGACCACCATTTTCACCATTTTGACAAATAAGGATGCAATCAAATCAGCCAAAGT

                510       520       530       540       550       560       570       580       590       600
        ....|....|....|....|....|....|....|....|....|....|....|....|....|....|....|....|....|....|....|....|
PipKuh  AGCCAAAGGAGTATCAAAACTATTTCATGAAAAACATAGGTCTTCAATCCACGAGGAAATGGAGAGGCTATTAGCGATTTGGATTAAGGACAGGCAGGTG
ErpCal  AGCCAAAGGAGTATCAAAACTATTTCATGAAAAACATAGGTCTTCAATCCACGAGGAAATGGAGAGGCTATTAGCGATTTGGATTAAGGACAGGCAGGTG

                610       620       630       640       650       660       670       680       690       700
        ....|....|....|....|....|....|....|....|....|....|....|....|....|....|....|....|....|....|....|....|
PipKuh  AAAGGTGACGTAACAACCCAAGATATTATCTGTCACAAAGCCAAGAGAATTTATGACGATCTAAAGAAAAACGTCCCTGGAAGTAGCAGCAATCAAGATA
ErpCal  AAAGGTGACGTAACAACCCAAGATATTATCTGTCACAAAGCCAAGAGAATTTATGACGATCTAAAGAAAAACGTCCCTGGAAGTAGCAGCAATCAAGATA

                710       720       730       740       750       760       770       780       790       800
        ....|....|....|....|....|....|....|....|....|....|....|....|....|....|....|....|....|....|....|....|
PipKuh  ATGAAGAAGAATTCAAAGCCAGCAGGGGGTGGTTTTTTAGATTTAAGAAAAGGTGTGGAATCCACAGCGTTACTATGCATGGTGAGGCTGGCAGTGCTGA
ErpCal  ATGAAGAAGAATTCAAAGCCAGCAGGGGGTGGTTTTTTAGATTTAAGAAAAGGTGTGGAATCCACAGCGTTACTATGCATGGTGAGGCTGGCAGTGCTGA


                810       820       830       840       850       860       870       880       890       900
        ....|....|....|....|....|....|....|....|....|....|....|....|....|....|....|....|....|....|....|....|
PipKuh  CAAGAAAGAAGCAGAAAAGTTCTCTATTAACTTTCAAAAATGTATTAAGGATGAAGGATACTGCCCACAACAAGTGTTCAATGCCGATGAAACGGGTCTT
ErpCal  CAAGAAAGAAGCAGAAAAGTTCTCTATTAACTTTCAAAAATGTATTAAGGATGAAGGATACTGCCCACAACAAGTGTTCAATGCCGATGAAACGGGTCTT

                910       920       930       940       950       960       970       980       990       1000
        ....|....|....|....|....|....|....|....|....|....|....|....|....|....|....|....|....|....|....|....|
PipKuh  TTCTGGAAAAGAATGCCGAGCAGAACCTTCATTACAAAAGAGGAGAAGAAATTGCCAGGACACAAAGCCATGAAGGACAGACTTACCCTTATGTTTTCGT
ErpCal  TTCTGGAAAAGAATGCCGAGCAGAACCTTCATTACAAAAGAGGAGAAGAAATTGCCAGGACACAAAGCCATGAAGGACAGACTTACCCTTATGTTTTCGT

                1010      1020      1030      1040      1050      1060      1070      1080      1090      1100
        ....|....|....|....|....|....|....|....|....|....|....|....|....|....|....|....|....|....|....|....|
PipKuh  CTAATGCCAGCGGAGACCTCAAGATCAAACCTCTATTGGTTTATCACTCTGAAAATCCAAGAATTTTCAAGAAAAATAACGTTATTAAGTCCAAACTGCC
ErpCal  CTAATGCCAGCGGAGACCTCAAGATCAAACCTCTATTGGTTTATCACTCTGAAAATCCCAGAATTTTCAAGAAAAATAACGTTATTAAGTCCAAACTGCC

                1110      1120      1130      1140      1150      1160      1170      1180      1190      1200
        ....|....|....|....|....|....|....|....|....|....|....|....|....|....|....|....|....|....|....|....|
PipKuh  CGTCCATTGGAAGTCCAATCAAAAAGCCTGGGTGACCCAAGTTATCTTCAACGAATGGATTCTGGAAACCTTTGCTCCTGCCGTGAAGAAATTCTTGCTG
ErpCal  CGTCCATTGGAAGTCCAATCAAAAAGCCTGGGTGACCCAAGTTATCTTCAACGAATGGATTCTGGAAACCTTTGCTCCTGCCGTGAAGAAATTCTTGCTG

                1210      1220      1230      1240      1250      1260      1270      1280      1290      1300
        ....|....|....|....|....|....|....|....|....|....|....|....|....|....|....|....|....|....|....|....|
PipKuh  GAAAAAGAACTGCCGCTCAAAGCCCTTCTGATACTTGACAATGCCCCTTCTCACCCAAAAGACCTAGAGGAAATATTGCAGGAAAATTATCCTTTTATCA
ErpCal  GAAAAAGAACTGCCGCTCAAAGCCCTTCTGATACTTGACAATGCCCCTTCTCACTCAAAAGACCTAGAGGAAATATTGCAGGAAAATTATCCTTTTATCA

                1310      1320      1330      1340      1350      1360      1370      1380      1390      1400
        ....|....|....|....|....|....|....|....|....|....|....|....|....|....|....|....|....|....|....|....|
PipKuh  AGGTGCAGTATTTGCCACCAAACACCACATCCATTCTTCAGCCAATGGATCAGCAAGTTATTGCGAACTTTAAAAAACTCTACACTAGAGCCCTCTTTAA
ErpCal  AGGTGCAGTATTTGCCACCAAACACCACATCCATTCTTCAGCCAATGGATCAGCAAGTTATTGCGAACTTTAAAAAACTCTACACTAGAGCCCTCTTTAA

                1410      1420      1430      1440      1450      1460      1470      1480      1490      1500
        ....|....|....|....|....|....|....|....|....|....|....|....|....|....|....|....|....|....|....|....|
PipKuh  TAAGGTGTTTGAAGAATGCGAGTTTGGTGGAGACAATATGACTGTCCGAAAGTTTTGGAAGGAGAAATTTGATGTCCTTATGGCAATACGACTTATACAG
ErpCal  TAAGGTGTTTGAAGAATGCGAGTTTGGTGGAGACAATATGACTGTCCGAAAGTTTTGGAAGGAGAAATTTGATGTCCTTATGGCAATACGACTTATACAG

                1510      1520      1530      1540      1550      1560      1570      1580      1590      1600
        ....|....|....|....|....|....|....|....|....|....|....|....|....|....|....|....|....|....|....|....|
PipKuh  AAGGCCTGGGAAGAAGTGTCACAAAGGACCCTCATTTCTGCTTGGAAGATGCTTGTGCCTTCGTGGACCCAGGAAGAAGCAGTAGTTGATGACACAGAAG
ErpCal  AAGGCCTGGGAAGAAGTGTCACAAAGGACCCTCATTTCTGCTTGGAAGATGCTTGTGCCTTCGTGGACCCAGGAAGAAGCAGTAGTTGATGACACAGAAG

                1610      1620      1630      1640      1650      1660      1670      1680      1690      1700
        ....|....|....|....|....|....|....|....|....|....|....|....|....|....|....|....|....|....|....|....|
PipKuh  TGGTGAAGGACATCATCACAGTGGCCCAAAGGTTGGAATTAGAGGTAGAGGAAGAGGATGTAGAGGAGCTTATTGAGGAACACGAAGAAGAGCTGACAAC
ErpCal  TGGTGAAGGACATCATCACAGTGGCCCAAAGGTTGGAATTAGAGGTAGAGGAAGAGGATGTAGAGGAGCTTATTGAGGAACACGAAGAAGAGCTGACAAC

                1710      1720      1730      1740      1750      1760      1770      1780      1790      1800
        ....|....|....|....|....|....|....|....|....|....|....|....|....|....|....|....|....|....|....|....|
PipKuh  TGAAGAGCTCCAAGCACTTCTGGTCCAGCAACAGGACAATGCTCAAAGGGAAGCGTCATCTGATAACGAGGAGCAACAATCAAACAATCAACCAATCCCA
ErpCal  TGAAGAGCTCCAAGCACTTCTGGTCCAGCAACAGGACAATGCTCAAAGGGAAGCGTCATCTGATAACGAGGAGCAACAATCAAACAATCAACCAATCCCA

                1810      1820      1830      1840      1850      1860      1870      1880      1890      1900
        ....|....|....|....|....|....|....|....|....|....|....|....|....|....|....|....|....|....|....|....|
PipKuh  ACTGCTGACATCAAGAACATCCTGGTCAAATGGAAAGCAGTTCAGGAGTTTACCAATGCCCACTATCCGGATTCAGCTGAAGCAAACAGGATCAACGATC
ErpCal  ACTGCTGACATCAAGAACATCCTGGTCAAATGGAAAGCAGTTCAGGAGTTTACCAATGCCCACTATCCGGATTCAGCTGAAGCAAACAGGATCAACGATC

                1910      1920      1930      1940      1950      1960      1970      1980      1990      2000
        ....|....|....|....|....|....|....|....|....|....|....|....|....|....|....|....|....|....|....|....|
PipKuh  TTTACTCCGATACTCTTGTCCGTTATTTCCGGCAGATGTTGGAGAAAAGAGAAAAACAAACGACTTTGGACAGGTTTTTCATGAAACCATCGGCCAAAAA
ErpCal  TTTACTCCGATACTCTTGTCCGTTATTTCCGGCAGATGTTGAAGAAAAGAGAAAAACAAACGACTTTGGACAGGTTTTTCATGAAACCATCGGCCAAAAA

                2010      2020      2030      2040      2050      2060      2070      2080      2090      2100
        ....|....|....|....|....|....|....|....|....|....|....|....|....|....|....|....|....|....|....|....|
PipKuh  GCAGAAAATGGATGAAGATGTGCAAGATTCGGTAGACTCTACTTAGTCTGTCTTAAAATTAAAAAAATGTGTGTTTTTTTTTTAAAAAAATTATGTTTTT
ErpCal  GCAGAAAATGGATGAAGATGTGCAAGATTCGGTAGACTCTACTTAGTCTGTCTTAAAATTAAAAAAATGTGTGTTTTTTT-AAAAAAAAATTATGTTTTT

                2110      2120      2130      2140      2150      2160      2170      2180      2190      2200
        ....|....|....|....|....|....|....|....|....|....|....|....|....|....|....|....|....|....|....|....|
PipKuh  AGATGTATCTAAATAAAAATAATAACAAAAAATTTATCTTTTTTTATGTCATCTTAGCATATTTTATGCTACAGAACGAATTATTTTTTTTAACATGTAT
ErpCal  AGATGTATCTAAATAAAAATAATAACAAAAAATTTATCTTTTTTTATGTCATCTTAGCATATTTTATGCTACAGAACGAATTATTTTTTTTAACATGTAT

                2210      2220      2230      2240      2250      2260      2270      2280      2290
        ....|....|....|....|....|....|....|....|....|....|....|....|....|....|....|....|....|....|.
PipKuh  TGTTATGGGAAAACGCGTTTCACATAACGAACTTTTCGCATAACAAACTTGCTCCTGGAACGAATTAAGTTCGTTGTGTGAGGCACCACTG
ErpCal  TGTTATGGGAAAACGCGTTTCACATAACGAACTTTTCGCATAACAAACTTGCTCCTGGAACGAATTAAGTTCGTTGTGTGAGGCACCACTG
```

**Fig. S1. Species-specific consensus sequence alignment for *Mariner2_pKuh* in the bat *Pipistrellus kuhlii* (PipKuh) and the African reedfish *Erpetoichthys calabaricus* (ErpCal).**

```
                 10        20        30        40        50        60        70        80        90       100
        ....|....|....|....|....|....|....|....|....|....|....|....|....|....|....|....|....|....|....|....|
AntPal  CTATATTTTCCAGCGTATAAGATGACTGGGCATATAAGACAACCCCTAACTTTTCCAGTTAAAATACAGAGTTTGGGATACACTTGCCCTATAAGAGGG-
EptFus  CCGTATTTTCCGGCGTATAAGACGACTGGGCGTATAAGACGACCCCCAACTTTTCCAGTTAAAATATAGAGTTTGGGATATACCCGCCCTATAAGATGA-
LasBor  CCGTATTTTCCGGCGTATAAGACGACTGGGCGTATAAGACGACCCCCAACTTTTCCAGTTAAAATATAGAGTTTGGGATATACTCGCCCTATAAGATGA-
LacAgi  CCGTATTTTCCGGCGTATAAGACGACTGGGCGTATAAGACGACCCCCAACTTTTCCAGTTAAAATATAGAGTTTGAGATATACTCGAC-CACA-GATTCT
ZooViv  CCGTATATTCCGGCGTATAAGACGACTGGGCGTATAAGACGACCCCCAACTTTTCCAGTTAAAATATAGAGTTTGGGATATACTCGCCGTATAAGAAATA

                110       120       130       140       150       160       170       180
        ....|....|....|....|....|....|....|....|....|....|....|....|....|....|....|....|..
AntPal  -CACCCAGCATATAAGACGACCCCCGACTTTTGAGAAGATTTTCCTGGGTTAAAAAGTCGTCTTATATGCCGGAAAATATGG
EptFus  -CACCCGGCGTATAAGACGACCCCCGACTTTTGAGAAGATTTTCCTGGGTTAAAAAGTCGTCTTATACGCCGGAAAATACGG
LasBor  -CACCCGGCGTATAAGACGACCCCCGACTTTTGAGAAGATTTTCCTGGGTTAAAAAGTCGTCTTATACGCCGGAAAATACGG
LacAgi  CCACCCGGCGTATAAGACGACCCCCGACTTTTGAGAAGATTTTCCTGGATTAAAAAGTAGTCTTATACGCCAGAATATACAG
ZooViv  CGACCCGGCGTATAAGACGACCCCCGACTTTTGAGAAGATTTTCCTGGGTTAAAAAGTAGTCTTATACGCAGGAATATACAG
```

**Fig. S2. Species-specific consensus sequence alignment for *nMariner1_Lbo* in the bats *Antrozous pallidus* (AntPal), *Eptesicus fuscus* (EptFus), and *Lasiurus borealis* (LasBor) and the lizards *Lacerta agilis* (LacAgi) and *Zootoca vivipara* (ZooViv).**

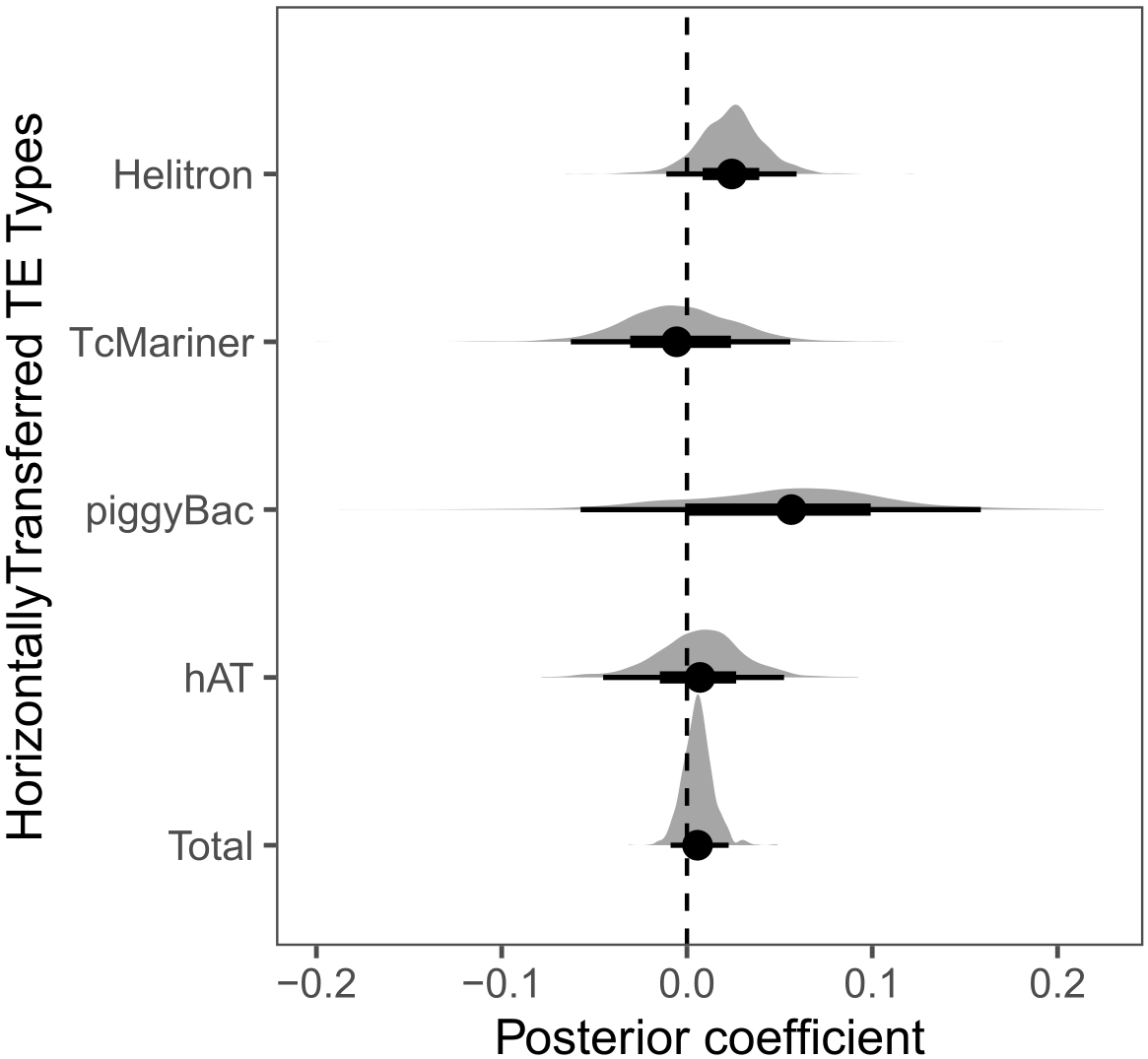
**Fig. S3. Posterior distributions of the regression coefficient of the proportion of species richness as a function of putative horizontally transferred TE diversity.** For each coefficient: black dots show median, thin lines show the 95% posterior probability, thick lines show the 66% posterior probability, and gray shows the posterior density of the estimates.

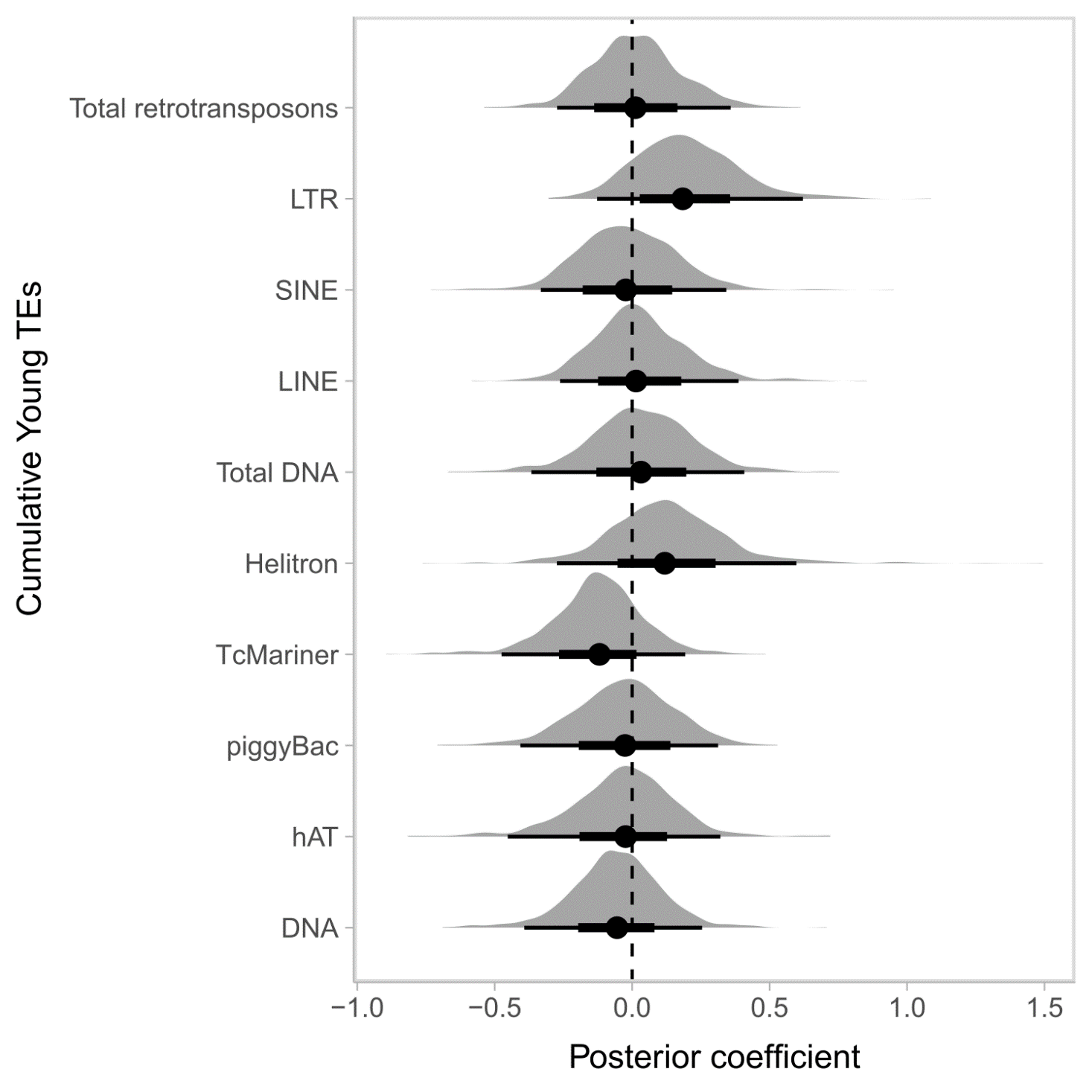
**Fig. S4. Posterior distributions of the regression coefficient of the proportion of species richness as a function of young (≤50 My) TE counts.** For each coefficient: black dots show median, thin lines show the 95% posterior probability, thick lines show the 66% posterior probability, and gray shows the posterior density of the estimates. “Total DNA” corresponds to total DNA transposons (all DNA transposons from hAT, piggyBac, Tc-Mariner, Helitron, and DNA categories). “DNA” corresponds to DNA transposons that have not been classified to a family level (such as hAT, Tc-Mariner, etc.).

**Supplemental Tables**

**Table S1. Eutherian mammal assemblies examined.**

**Table S2. TE content for each eutherian mammal species examined.** RT, Retrotransposon; DNA, DNA transposon; RC, Rolling-circle transposon.

**Table S3. Calculated neutral mutation rates of 251 mammals (37 bats, 214 other eutherian mammals).** Branch lengths taken from Foley et al. (2022); divergence times taken from TimeTree, accessed 2 February 2023.

**Table S4. Identities of putative horizontally transferred TEs.**

**Table S5. Hit counts of putative horizontal TE transfers and the species involved.**

**Table S6. Branch assignments of putative horizontally transferred transposons.**

**Table S7. Estimated ages for putative horizontal transfer events with non-chiropteran matches.**

**Table S8. Input data for model posterior distributions of group category on horizontal transfer event counts (Fig. 3).**

**Table S9. Calculated rates of horizontal transfer events in Chiroptera.**

**Table S10. Branch numbering and divergence times based on TimeTree. Non-conflicting combinations of median and average divergence times were used in the phylogenetic tree in Fig. 4.**

**Table S11. Summary of posterior distributions of the regression coefficient of the proportion of species richness as a function of the modeled TE diversity counts.** ESS, Estimated sampling size; HPD, high probability density interval; l, lower; PSRF, potential scale reduction factor; u, upper.

| Horizontally transferred TE | Estimate | l-95% HPD | u-95% HPD | PSRF | ESS |
| --- | --- | --- | --- | --- | --- |
| Total | 0.01 | -0.01 | 0.02 | 1.00 | 1589 |
| hAT | 0.01 | -0.05 | 0.05 | 1.00 | 1737 |
| piggyBac | 0.05 | -0.06 | 0.16 | 1.00 | 1603 |
| Tc-Mariner | 0.00 | -0.06 | 0.06 | 1.00 | 1295 |
| Helitron | 0.02 | -0.01 | 0.06 | 1.00 | 1597 |

**Table S12. Summary of posterior distributions of the regression coefficient of the proportion of species richness as a function of the log10-transformed, scaled TE insertion counts.** ESS, Estimated sampling size; HPD, high probability density interval; l, lower; PSRF, potential scale reduction factor; u, upper.

| Cumulative TE | Estimate | l-95% HPD | u-95% HPD | PSRF | ESS |
| --- | --- | --- | --- | --- | --- |
| DNA | -0.06 | -0.39 | 0.25 | 1.00 | 914 |
| hAT | -0.03 | -0.44 | 0.34 | 1.00 | 1048 |
| piggyBac | -0.04 | -0.41 | 0.29 | 1.00 | 1717 |
| Tc-Mariner | -0.13 | -0.51 | 0.21 | 1.00 | 4117 |
| Helitron | 0.13 | -0.27 | 0.60 | 1.00 | 3878 |
| Total DNA transposons | 0.04 | -0.35 | 0.43 | 1.00 | 1275 |
| LINE | 0.03 | -0.26 | 0.39 | 1.00 | 1189 |
| SINE | -0.01 | -0.34 | 0.33 | 1.00 | 1574 |
| LTR | 0.19 | -0.14 | 0.59 | 1.00 | 1439 |
| Total Retrotransposons | 0.02 | -0.29 | 0.38 | 1.00 | 1392 |

**Table S13. Assembly statistics for available bat species.**
